# ZFP36 RNA-binding proteins restrain T-cell activation and anti-viral immunity

**DOI:** 10.1101/247668

**Authors:** Michael J. Moore, Nathalie E. Blachere, John J. Fak, Christopher Y. Park, Kirsty Sawicka, Salina Parveen, Ilana Zucker-Scharff, Bruno Moltedo, Alexander Y. Rudensky, Robert B. Darnell

**Affiliations:** Laboratory of Molecular Neuro-Oncology, and Howard Hughes Medical Institute, The Rockefeller University, New York, NY, USA.; New York Genome Center, 101 Avenue of the Americas, New York, NY, USA; Ludwig Center at Memorial Sloan Kettering Cancer Center and Howard Hughes Medical Institute, New York, NY, USA; Present Affiliation: Regeneron Pharmaceuticals, Tarrytown, NY, USA

## Abstract

Dynamic post-transcriptional control of RNA expression by RNA-binding proteins (RBPs) is critical during immune response. ZFP36 RBPs are prominent inflammatory regulators linked to autoimmunity and cancer, but functions in adaptive immunity are less clear. We used HITS-CLIP to define ZFP36 targets in T-cells, which revealed unanticipated actions in regulating T-cell activation, proliferation, and effector functions. Transcriptome and ribosome profiling showed that ZFP36 represses mRNA target abundance and translation, notably through a novel class of AU-rich sites in coding sequence. Functional studies revealed that ZFP36 regulates early T-cell activation kinetics in a cell autonomous manner, by attenuating activation marker expression, limiting T-cell expansion, and promoting apoptosis. Strikingly, loss of ZFP36 in vivo accelerated T-cell responses to acute viral infection, and enhanced anti-viral immunity. These findings uncover a critical role for ZFP36 RBPs in restraining T-cell expansion and effector functions, and suggest ZFP36 inhibition as a novel strategy to enhance immune-based therapies.

## Introduction

Immune responses require precise, dynamic gene regulation that must activate rapidly as threats rise, and resolve efficiently as they clear. Post-transcriptional control of mRNA abundance and expression by RNA binding proteins (RBPs) is a key layer of this response that can enact rapid, signal-responsive changes (Kafasla et al. 2014; Hao and Baltimore 2009), but knowledge of specific functional roles for dedicated RBPs remains limited.

AU-rich elements (AREs) in mRNA 3’-untranslated regions (3’-UTR) facilitate post-transcriptional control of many immune functions, including cytokine expression, signal transduction, and immediate-early transcriptional response (Chen and Shyu 1994; Shaw and Kamen 1986; Caput et al. 1986). Many RBPs bind AREs, with diverse ensuing effects on RNA turnover, translation, and localization (Stoecklin and Mühlemann 2013; Tiedje et al. 2012; Roretz et al. 2011). The Zinc finger binding protein 36 (ZFP36) family of proteins are prototypical ARE-binding factors with distinctive, activation-dependent expression in hematopoietic cell lineages (Raghavan et al. 2001; Carballo et al. 1998). The family includes three somatic paralogs: ZFP36 (a.k.a Tristetraprolin, TTP), ZFP36L1 (a.k.a. butyrate responsive factor 1, BRF1), and ZFP36L2 (a.k.a BRF2) with highly homologous CCCH zinc-finger RNA binding domains (Blackshear 2002).

In many contexts, the archetypal paralog ZFP36 de-stabilizes target mRNAs by binding 3’-UTR AREs and recruiting deadenylation and degradation factors (Brooks and Blackshear 2013; Lykke-Andersen and Wagner 2005). More recently, evidence has begun to emerge for roles in translation (Tao and Gao 2015; Tiedje et al. 2012) but in vivo function and context have not been established. While many aspects remain unsettled, ZFP36 is clearly critical for immune function, as its loss causes a systemic inflammatory disease in mice (Taylor et al. 1996). A key feature of this syndrome is aberrant stabilization and over-expression of *Tnf* in myeloid cells, particularly macrophages (Carballo et al. 1998), where UV cross-linking and immunoprecipitation (CLIP) analyses have further supported direct regulation (Tiedje et al. 2016; Sedlyarov et al. 2016). However, this role does not fully account for ZFP36 function in vivo, as underscored by reports that myeloid-specific deletions of *Zfp36* do not recapitulate spontaneous autoimmunity (Qiu et al. 2012; Kratochvill et al. 2011).

Increasing evidence points to important functions for ZFP36 proteins in adaptive immunity. Dual ablation of paralogs *Zfp36l1* and *Zfp36l2* in T-cells arrests thymopoeisis at the double-negative stage, and causes lethal lymphoma linked to *Notch1* dysregulation (Hodson et al. 2010). This role in restraining aberrant proliferation was later extended to B-cell development and lymphoma (Galloway et al. 2016; Rounbehler et al. 2012), but the severe phenotype precluded analysis of ZFP36 family function in mature T-cells. Consistent with such a function, in vitro studies suggest ZPF36 regulates the expression of T-cell-derived cytokines, including IL-2, IFN-γ and IL-17, that mediate lymphocyte homeostasis, microbial response, and inflammation (Lee et al. 2012; Ogilvie et al. 2009; 2005). The landscape of ZFP36 targets beyond these limited cases in T-cells is unknown, but will be the key to understanding its emerging roles in inflammation, autoimmunity, and malignant cell growth (Patial and Blackshear 2016).

To determine ZFP36 functions in T-cells, we employed high-throughput sequencing of UV-cross-linking and immunoprecipitation (HITS-CLIP) to generate a definitive set of ZFP36 RNA targets. HITS-CLIP utilizes in vivo UV-cross-linking to induce covalent bonds between RBPs and target RNAs, allowing stringent immunopurification and thus rigorous identification of direct binding events (Licatalosi et al. 2008; Ule et al. 2003). These new ZFP36 RNA binding maps pointed to roles in regulating T-cell activation kinetics and proliferation, a function confirmed in extensive functional assays, and in vivo studies demonstrating a critical role in anti-viral immunity. Our results illuminate novel functions for ZFP36 in adaptive immunity, laying groundwork for understanding and modulating its activity in disease.

## Results

### ZFP36 dynamics during T-cell activation

ZFP36 expression is induced upon T-cell activation (Raghavan et al. 2001). We examined its precise kinetics following activation of primary mouse CD4+ T-cells by Western analysis with custom ZFP36 antisera. Protein levels were induced rapidly after activation, peaking at ~4 hours, tapered gradually through 72 hours, and were re-induced by re-stimulation 3 days post-activation (Figure 1A). ZFP36 expression depended on both TCR stimulation, provided by anti-CD3, and co-stimulation, provided by co-cultured dendritic cells (DCs) (Figure 1B). A similar pattern of transient ZFP36 induction occurred in activated CD8+ T-cells (Figure S1A).

Western blots showed multiple bands at ~40-50 kD, reflecting previously reported phospho-isoforms (Qiu et al. 2012). We also examined ZFP36L1 and ZFP36L2, as the antisera epitope is conserved across paralogs, and cross-reactivity was confirmed using recombinant constructs (Figure S1B). Analysis of *Zfp36* KO T-cell lysates with this antisera (henceforth, anti-pan-ZFP36) showed ~50% reduced signal compared to WT (Figure S1C). As ZFP36 and ZFP36L1 are highly homologous and of similar size, we interpret the signal in *Zfp36* KO cells as ZFP36L1. Accordingly, paralog-specific antibodies confirmed that both ZFP36 and ZFP36L1 were induced by T-cell activation (Figure S1D). ZFP36L2, expected to run at ~62 kD, was not detected in these analyses. These results demonstrate activation-dependent expression of ZFP36 and ZFP36L1 in T-cells, and suggest *Zfp36* KO T-cells have partial loss of pan ZFP36 activity.

**Figure 1.**
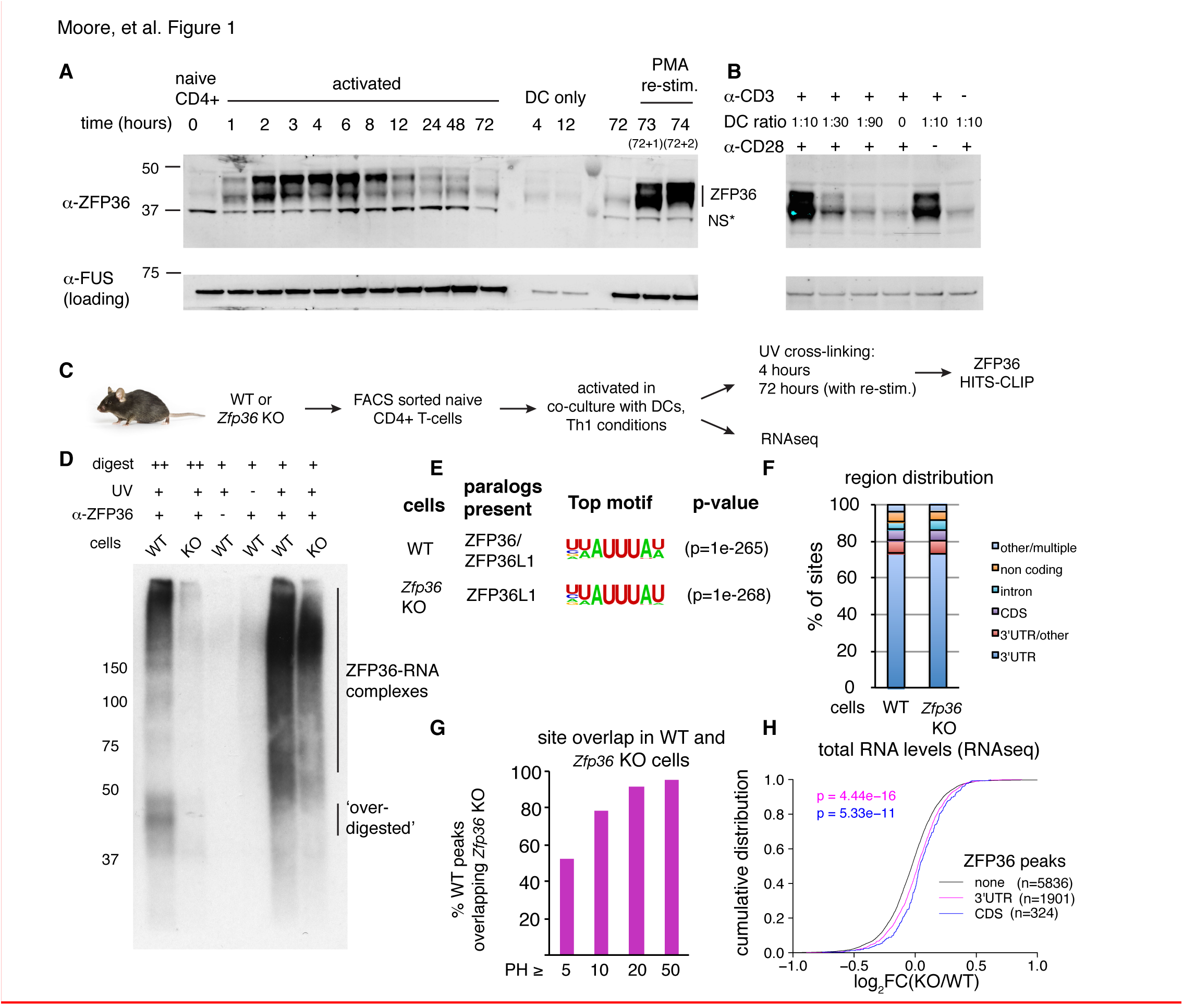
HITS-CLIP as a transcriptome-wide screen for ZFP36 function in T-cells. **(A)** Immunoblots with pan-ZFP36 antisera after activation of naïve CD4+ T-cells in DC co-cultures, and with re-stimulation at day 3. Antibody and MW markers are shown on the left. NS* indicates a non-specific band. **(B)** Immunoblotting with pan-ZFP36 antisera 4 hours after activation of naïve CD4+ T-cells, testing dependence on TCR stimulation (α-CD3), and co-stimulation (DCs or α-CD28). **(C)** ZFP36 HITS-CLIP design. **(D)** Representative autoradiogram of ZFP36 CLIP from activated CD4+ T-cells using pan-ZFP36 antisera, with pre-immune and no-UV controls. RNP signal in *Zfp36* KO cells is due to ZFP36L1. **(E)** The most enriched binding motifs and **(F)** annotation of binding sites from WT and *Zfp36* KO cells. **(G)** Overlap of binding sites in WT and *Zfp36* KO cells, stratified by peak height (PH).**(H)** RNAseq in WT and *Zfp36* KO CD4+ T-cells activated underTh1 conditions for 4 hours. Log2-transformed fold-changes (KO/WT) are plotted as a cumulative distribution function (CDF), for mRNAs with 3’UTR, CDS, or no significant ZFP36 HITS-CLIP sites. Numbers of mRNAs in each category (n) and p-values from two-tailed Kolmogorov-Smirnov (KS) tests are shown. See also Figures S1-S2.

**Supplementary Figure S1.**
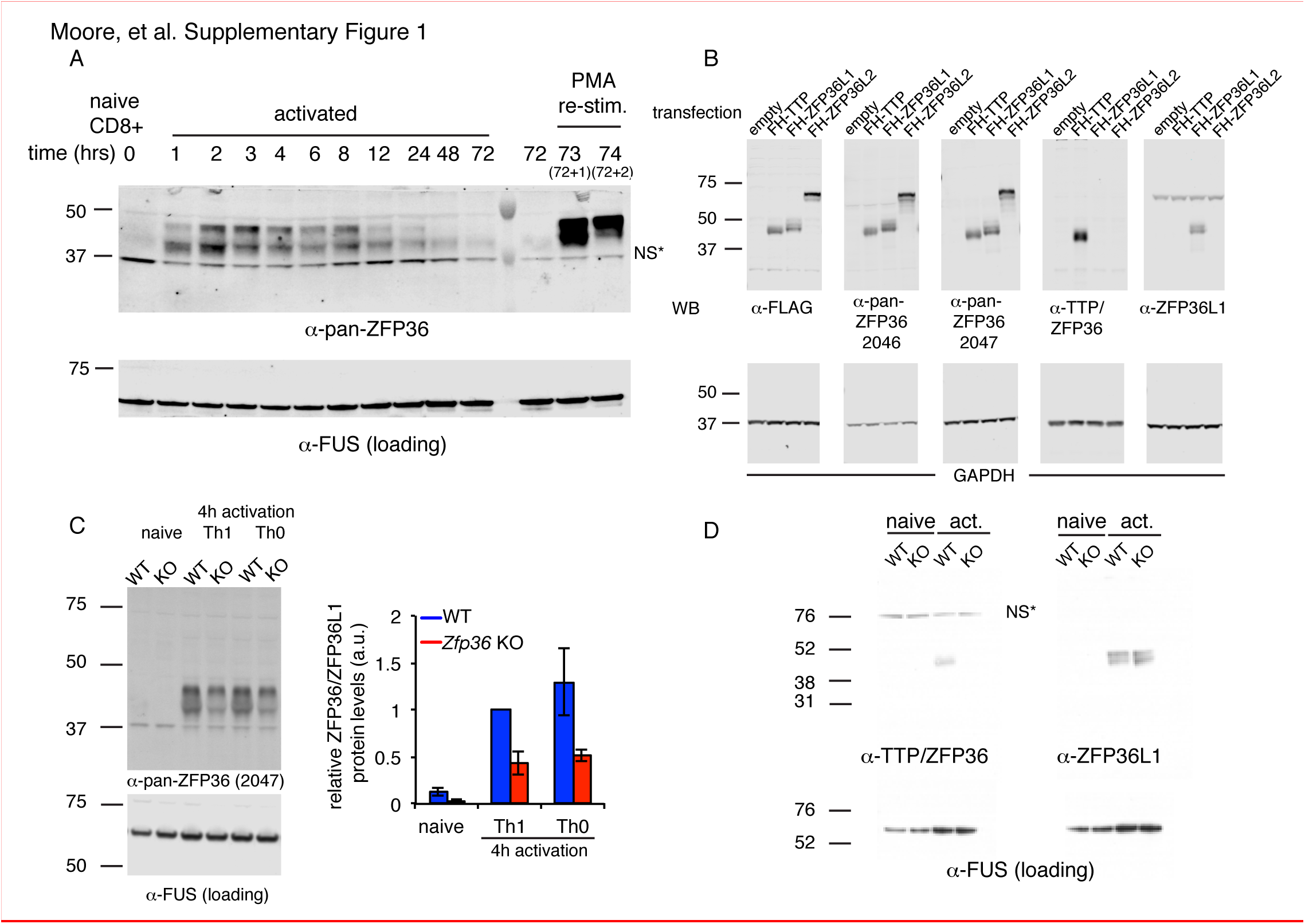
ZFP36 paralog expression in T-cells. **(A)** ZFP36 expression was tracked in atime course of CD8+ T-cell activation using pan-ZFP36 antisera. Cells re-stimulated with PMA/ionomycin after 3 days in culture were also analyzed. **(B)** 293T cells were transfected with indicated constructs expressing FLAG-HA (FH) tagged ZFP36 paralogs. Lysates were analyzed by immunoblotting with antibodies indicated below panels: anti-FLAG; two custom antisera (RF2046 and RF2047) raised against a C-terminal peptide of mouse ZFP36; commercial anti-ZFP36; and commercial anti-ZFP36L1. NS* indicates a presumably cross-reacting band. **(C)** Lysates from naïve, Th1-, or Th0-activated WT and *Zfp36* KO CD4+ T-cells (4 hours) were analyzed by immunoblotting with pan-anti-ZFP36. A representative blot is shown (left), with quantification across three independent experiments. **(D)** Lysates from naïve or Th1-activated WT and *Zfp36* KO CD4+ T-cells were analyzed by immunoblotting with ZFP36- and ZFP36L1-specific antibodies.

### Transcriptome-wide identification of ZFP36 target RNAs in CD4+ T-cells

The striking pattern of ZFP36/L1 expression in T-cells led us to develop ZFP36 HITS-CLIP as a screen for its biological function (Figure 1C–F; Figure S2). Notably, ZFP36/L1 RNPs isolated by CLIP from activated CD4+ T-cells sera exhibited high molecular weight (MW) complexes resistant to detergent, heat, and RNAse, consistent with a pattern previously observed in ZFP36 CLIP in macrophages (Figure 1D, Figure S2A; (Sedlyarov et al. 2016)). This RNP signal pattern was dependent on UV irradiation, and was observed with two different anti-pan-ZFP36 but neither pre-immune sera.

**Supplementary Figure S2.**
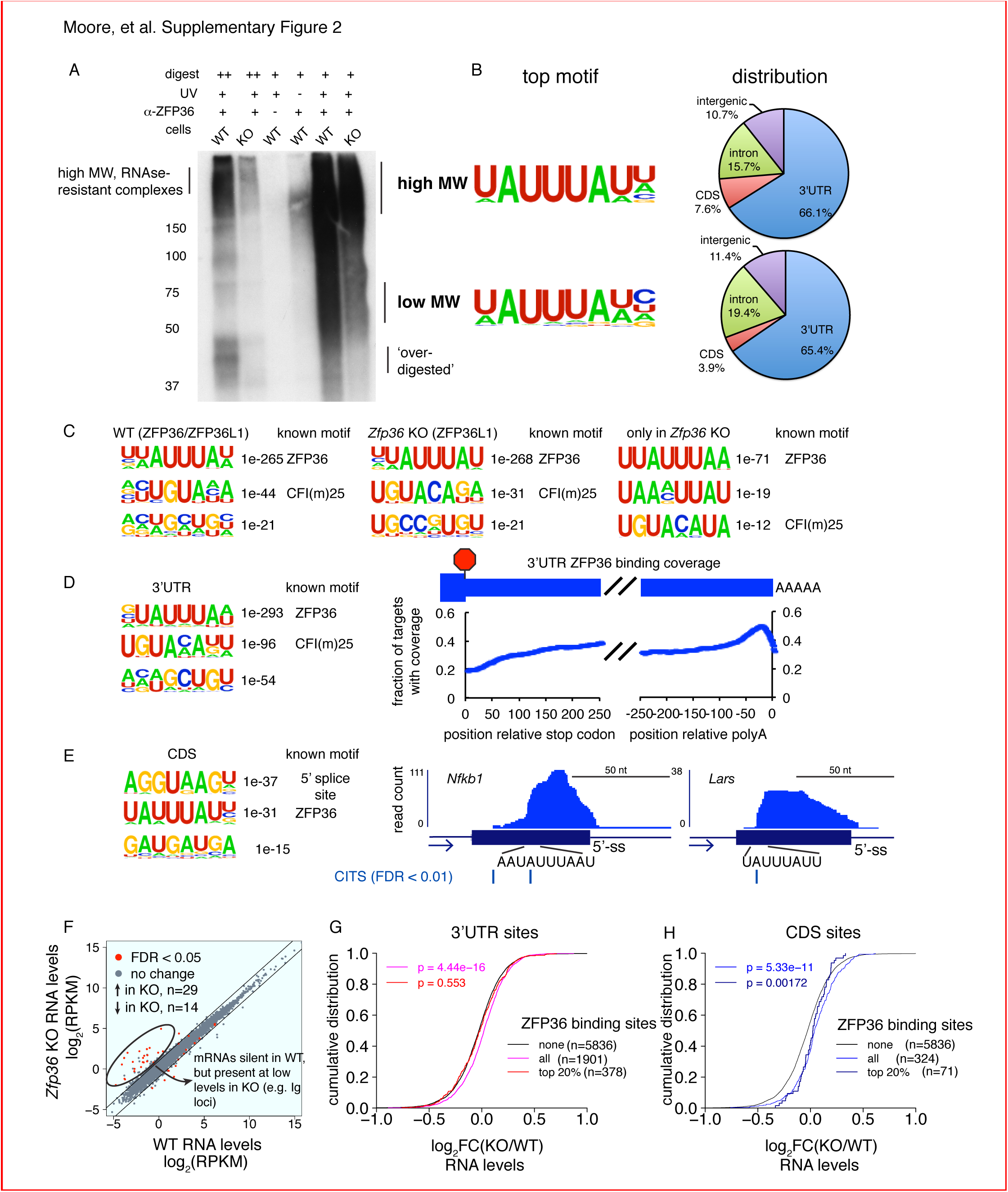
ZFP36 HITS-CLIP. **(A)** ZFP36 HITS-CLIP autoradiogram with a second pan-ZFP36 antisera (RF2046) is shown, with similar results to RF2047 (Figure 1D). **(B)** High MW and low MW ZFP36-RNA complexes were analyzed separately, yielding sites with very similar enriched motifs and transcript distribution. **(C)** Top three enriched motifs from HITS-CLIP sites in WT cells (left), *Zfp36* KO cells (middle), and sites only identified in *Zfp36* KO cells (right). Correspondence to previously known motifs is shown. **(D)** Motif analysis of ZFP36 HITS-CLIP sites from WT cells in 3’UTR (left), and the position of ZFP36 binding sites in 3’UTR depicted as a coverage map (right). **(E)** Motif analysis of ZFP36 HITS-CLIP sites from WT cells in CDS (left), and representative CDS binding sites visualized in UCSC Genome Browser. CLIP reads supporting CDS binding sites span exon-intron boundaries, consistent with 5’-ss motif enrichment. Cross-link-induced truncations (CITS) are shown, confirming that binding occurs within the coding exon. **(F)** Transcriptome profiling data plotted as RPKM in *Zfp36* KO cells versus RPKM in WT cells. Few large changes were observed, with the exception of highlighted mRNAs (e.g. Ig mRNAs) that were silent in WT cells, but show expression in KO cells. The absence of CLIP binding sites in this subset suggests secondary effects. (**G)** Transcriptome profiling results are depicted as a CDF as in Figure 1, further stratified by the magnitude of ZFP36 binding in 3’UTR. 3’UTR target mRNAs overall (violet) show a significant shift relative to non-targets (black), but the top 20% of sites defined by peak height (red) show no significant shift. Numbers of mRNAs in each category (n) are indicated, and p-values show results of Kolmogorov-Smirnov (KS) test. **(H)** Analysis as in **(H)** for mRNAs with ZFP36 binding in CDS.

Given prior evidence that ZFP36 regulates T-helper type-1 (Th1) cytokines (e.g. TNF-α and IFN-γ), we next generated a comprehensive map of ZFP36/L1 binding sites by HITS-CLIP using anti-pan-ZFP36 in WT CD4+ T-cells, activated for 4 hours under Th1-polarizing conditions (Ogilvie et al. 2009; Carballo et al. 1998). 5132 robust binding sites were defined, requiring a peak height (PH) > 5, and support from at least 3 (of 5 total) biological replicates and two different pan-ZFP36 antisera (Table S1A; (Shah et al. 2016; Moore et al. 2014)). Consistent with identification of bona fide ZFP36/L1 binding events, HITS-CLIP recovered the known AU-rich ZFP36 consensus motif at high significance, along with reported binding sites in *Tnf*, *Ifng* and other targets (Figure 1E; Table S1A; (Brewer et al. 2004)). Globally, ZFP36/L1 sites confirmed a preponderance of 3’-UTR binding (>75%), and showed substantial binding in coding sequence (CDS; ~6.5%) and introns (5.4%) (Figure 1F). Separate analysis of low and high MW RNP complexes showed similar transcript localization and motif enrichment, all consistent with ZFP36 binding (Figure S2B). This analysis indicates the presence of large, stable ZFP36 complexes in vivo, consistent with stable multimers (Cao et al. 2003; Cao 2004). Subsequently, CLIP reads from different MW regions were pooled to maximize dataset depth.

To examine possible paralog specificity, we also mapped ZFP36L1 sites by HITS-CLIP in *Zfp36* KO CD4+ T-cells under identical conditions. As in Western analysis (Figure S1C–D), *Zfp36* KO samples showed reduced but significant CLIP signal compared to WT (Figure 1D, Figure S2A), representing ZFP36L1-RNA complexes. Sites in WT and *Zfp36* KO cells showed very similar enriched motifs and transcript localizations, indicating that ZFP36 and ZFP36L1 have similar binding profiles in vivo (Figures 1E–F and S2C, Table S1B). Majorities of robust sites (53%) and target mRNAs (66%) identified in WT cells were found independently in *Zfp36* KO cells, and site overlap was far greater (>90%) for peaks of increasing magnitude (Figure 1G). A subset of potential ZFP36L1-specific sites was identified only in *Zfp36* KO cells (n=675; Table S1C), although these showed similar features to ZFP36 sites overall (Figure S2C; third panel). Thus, these analyses do not exclude paralog specificity at some sites, but indicate broadly redundant in vivo RNA binding for ZFP36 and ZFP36L1 reflecting their high homology.

Secondary enriched motifs revealed additional properties of ZFP36/L1 target sites. The second top motif resembled the known recognition sequence for polyadenylation factor CFI(m)25 (Venkataraman et al. 2005). Accordingly, ZFP36 binding in 3’UTRs was most concentrated in the vicinity of expected polyA sites, ~50 nucleotides before transcript ends (Figure S2D). Analysis of CDS-specific binding revealed the AU-rich ZFP36 motif, along with strong enrichment of the 5’ splice site (5’-ss) consensus (Figure S2E). Cross-link-induced truncations (CITS) clarified that CDS peaks are centered within coding exons, but supporting CLIP reads often spanned the exon-intron boundary. Thus, at least a subset of CDS binding by ZFP36 occurs prior to pre-mRNA splicing in the nucleus.

### ZFP36 represses target mRNA abundance and translation during T-cell activation

We next employed RNA profiling strategies to determine the functional effects of ZFP36 binding. RNAseq analysis in WT and *Zfp36* KO CD4+ T-cells activated under conditions identical to our HITS-CLIP analyses uncovered two effects. First, a small number of mRNAs that were silent in WT cells, including immunoglobulin loci, were detected at low levels in KO cells (Figure S2F). These mRNAs lacked evidence of ZFP36 binding, suggesting secondary effects, such as dysregulated chromatin or transcriptional silencing at these loci. Second, a global analysis revealed that ZFP36 binding in 3’UTR (p=4.44x10^-16^; Kolmogorov-Smirnoff [KS]) and CDS (p=5.33x10^-11^) correlated to subtle but highly significant shifts toward greater mRNA abundance in *Zfp36* KO cells, relative to mRNAs with no binding sites (Figure 1H). This correlation was not observed for mRNAs with binding exclusively in introns, and we did not find evidence in these data that ZFP36 binding correlated with altered usage of proximal splice or polyA sites (not shown).

The overall trend in transcriptome profiling is consistent with evidence that ZFP36 represses RNA abundance (Lykke-Andersen and Wagner 2005). However, stratification of sites by the magnitude of ZFP36 binding allowed resolution of potentially complex effects. For 3’UTR binding, ZFP36 targets overall showed a significant shift in abundance, but mRNAs containing the top 20% most robust sites (ranked by peak height [PH], see Methods) showed no significant effect. Thus, a higher degree of bindinconstruct showed increased g correlated with less effect on RNA abundance in the absence of ZFP36 (Figure S2G). This trend was not observed for CDS sites, where the top 20% showed a similar shift to sites overall. Thus, our analyses show a trend of negative regulation of RNA abundance in this context, but with blunted effects for highly robust binding sites in 3’UTR (see Discussion). The same pattern was also observed when considering a more stringently defined of sites set (n=2178) overlapping statistically robust CITS.

We further investigated the effects of ZFP36 regulation for HITS-CLIP targets with highly robust 3’UTR binding in T-cells. Activation marker CD69, apoptosis regulator BCL2, and effector cytokines TNF and IFNG showed significantly increased protein levels in *Zfp36* KO versus WT activated T-cells. Of these, only *Bcl2* showed increased mRNA abundance (Figure 2A). *Tnf,Ifng*, and *Cd69* were all among the top 20% of targets as defined by CLIP binding magnitude (PH), thus supporting the trend observed in our global analyses that some highly robust binding targets show little regulation at the level of mRNA abundance in this context. The effects on protein level in the absence of changes in mRNA abundance of these targets suggest effects on translation. We tested this possibility by constructing Acgfp1 fluorescent reporters with an intact *Ifng* 3’UTR (WT-UTR), or with the CLIP-defined ZFP36 binding site deleted (Δ-UTR; Figure 2B). In 293 cells, ZFP36 repressed RNA levels for the WT-UTR reporter ~1.5-fold, but not the Δ-UTR construct (Figure 2C), extending prior data that ZFP36 over-expression represses *Ifng* mRNA levels in vitro (Ogilvie et al. 2009). Critically, ZFP36 exerted much greater ~5-fold repression of reporter protein levels (Figure 2C, right panel). Of note, the Δ-UTR construct showed increased protein levels both in the presence and absence of ZFP36. In addition, ZFP36-mediated repression of protein from WT-UTR (5-fold) was only partially relieved for Δ-UTR (1.7-fold). Thus, additional factors likely regulate this site in 293 cells. Nevertheless, the greater repressive effect of ZFP36 on target protein versus RNA levels is further consistent with regulation of translation, in addition to RNA stability.

**Figure 2.**
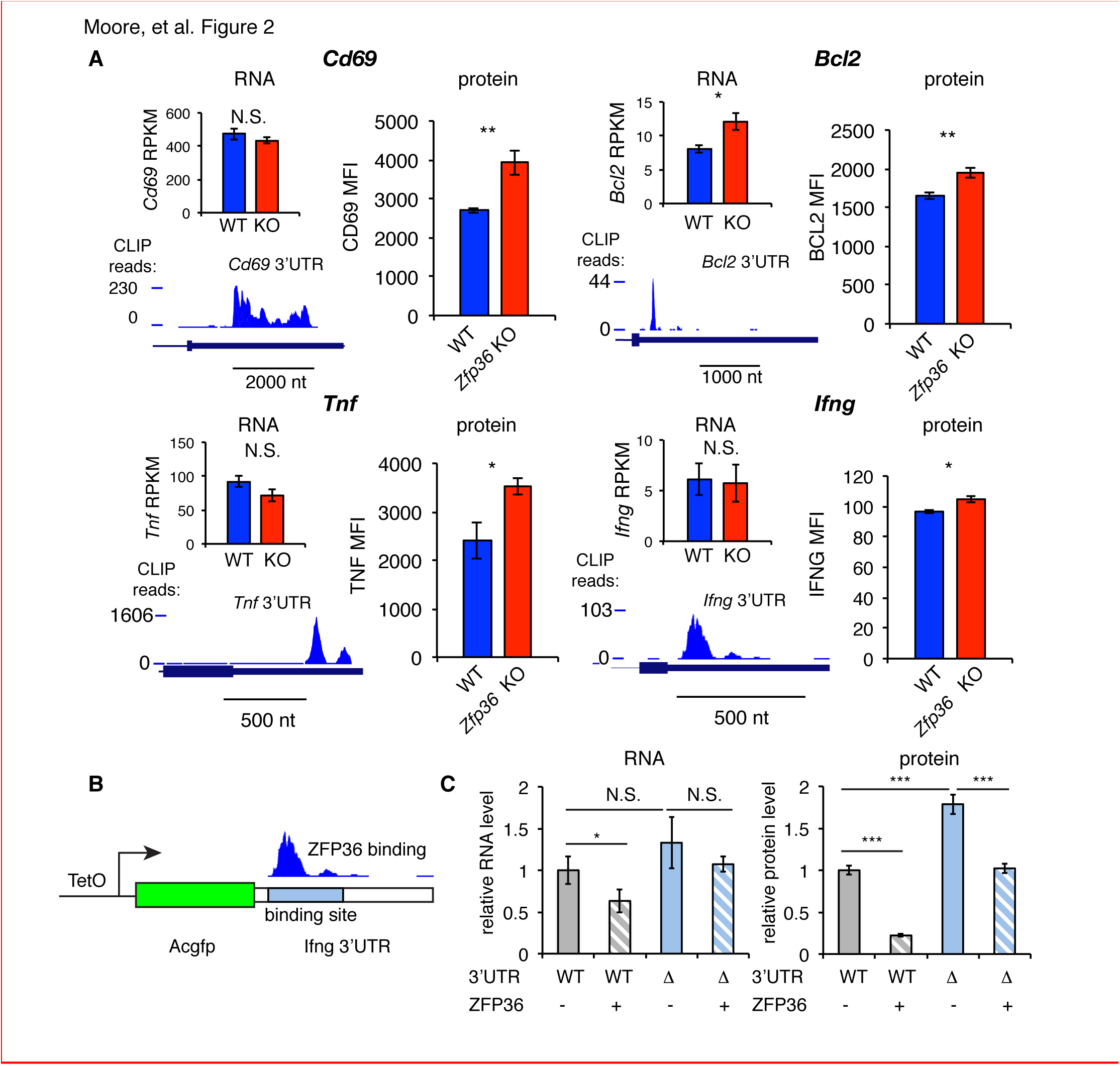
ZFP36 regulates target protein levels in T-cells. **(A)** Levels of mRNA and protein in *Zfp36* KO and WT T-cells and ZFP36 CLIP tracks are shown 4 hours post-activation for targets with robust 3’UTR ZFP36 binding. RNA values are mean RPKM +/-S.E.M. of 4 biological replicates. Protein values are mean fluorescence intensities (MFI) +/- S.E.M. for 3-4 mice per condition. **(B)** A reporter assay using AcGFP1 under a Tet-inducible promoter, with the WT *Ifng* 3’UTR (WT-UTR) or one lacking the ZFP36 binding site (-UTR). **(C)** WT-UTR or-UTR reporters were co-transfected into 293 cells with *Zfp36* (+) or vector alone (-). 24 hours post-transfection, reporters were induced with doxycycline for 4 hours, then RNA and protein levels were measured by RT-qPCR and flow cytometry, respectively. Values are mean +/- S.D. of 4 biological replicates in each condition. Results of two-tailed t-tests: * = p<0.05; ** =p<0.01; *** =p<0.001.

To directly test a role for ZFP36 in translational regulation in T-cells, we performed ribosome profiling of WT and *Zfp36* KO CD4+ T-cells activated for 4 hours under Th1 conditions (Figures 3A and S3A; (Ingolia et al. 2009)). We observed robust ribosome association of *Zfp36* mRNA in WT cells that was lost completely downstream of the engineered gene disruption in *Zfp36* KO cells (Figure S3B), consistent with accurate identification of translating mRNAs. Globally, there was a subtle but significant shift toward greater ribosome association in *Zfp36* KO cells for mRNAs bound by ZFP36 in 3’UTR (p=1.04x10^-11^; KS) or CDS (p=1.58x10^-11^), relative to mRNAs with no ZFP36 binding (Figure 3B). These shifts mirror those for global RNA abundance, with two notable exceptions. First, mRNAs with ZFP36 binding in CDS showed a significantly larger shift in ribosome association than those with 3’UTR binding (Figure 3C). Second, in contrast to effects on RNA abundance, the top 20% most robust ZFP36 binding sites in 3’UTR showed similar effects on ribosome association to sites overall (Figure S3C). Thus, ZFP36 target mRNAs show increased ribosome association in *Zfp36* KO cells.

**Figure 3.**
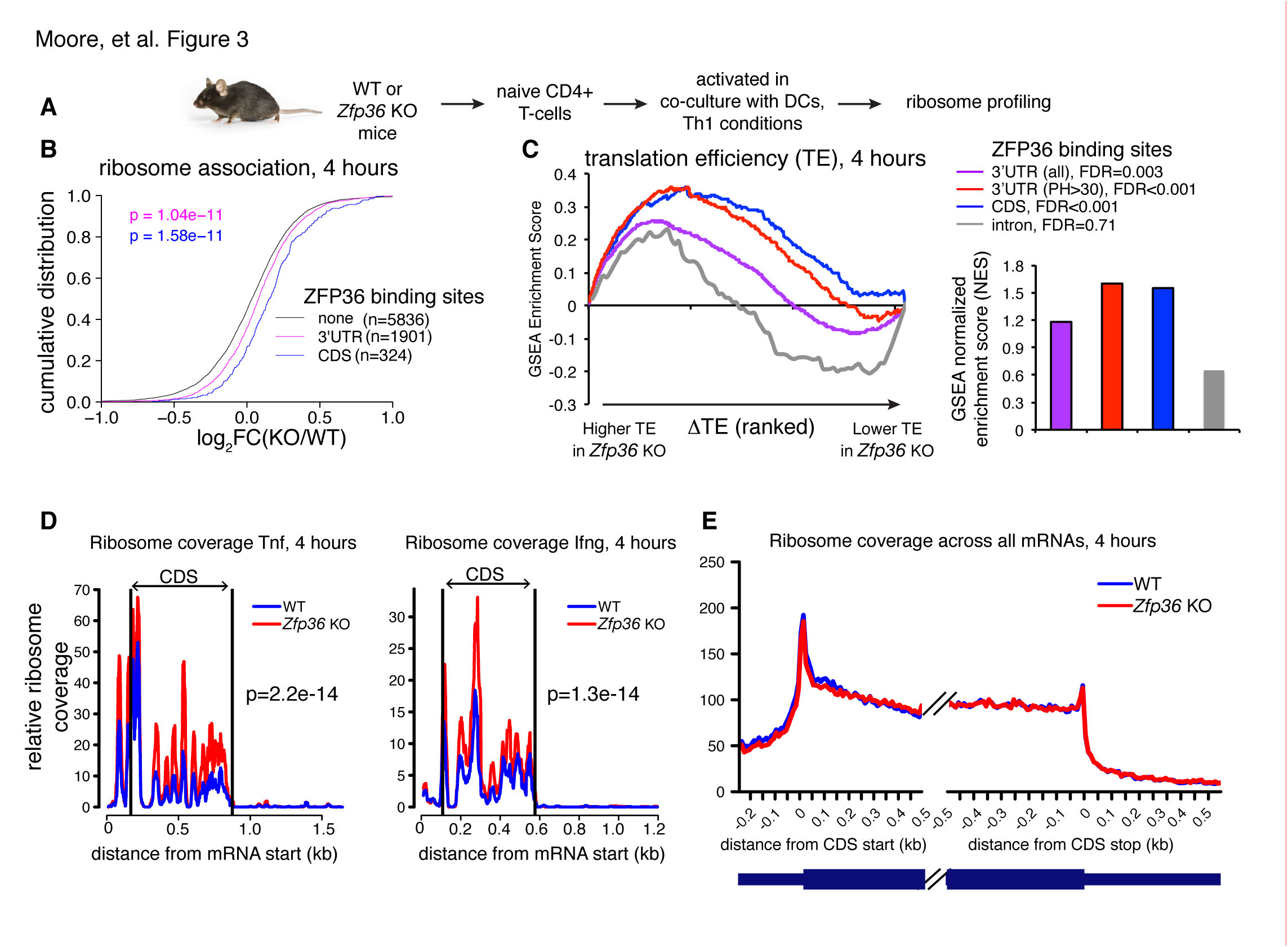
ZFP36 regulates target ribosome association. **(A)** Ribosome profiling of *Zfp36* KO and WT CD4+ T-cells. **(B)** Changes in ribosome association between *Zfp36* KO and WT cells plotted as a CDF. **(C)** Change in translation efficiency (ΔTE) between *Zfp36* KO and WT was calculated as a delta between log2(KO/WT) from ribosome profiling and RNAseq. The distribution of ZFP36 targets in mRNAs ranks by TE is shown (left), along with normalized enrichment scores and FDRs from GSEA (right). **(D)** Normalized coverage of ribosome profiling reads for *Tnf* and *Ifng* mRNAs in *Zfp36* KO and WT cells, with p-values from binomial tests. **(E)** Normalized coverage of ribosome profiling reads across all mRNAs for *Zfp36* KO and WT cells. See also Figure S3.

**Supplementary Figure S3.**
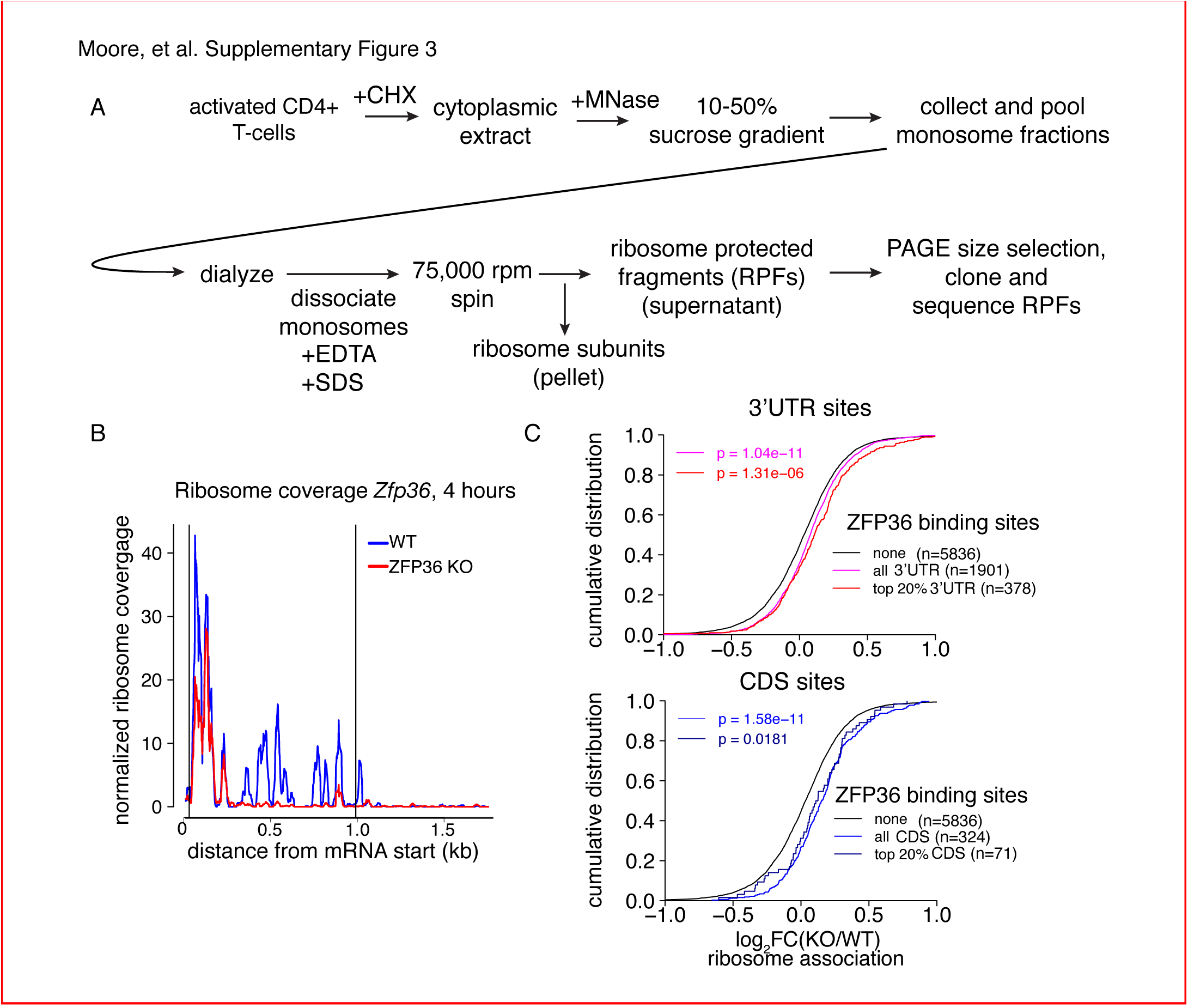
Analysis of ZFP36 translational control by ribosome profiling. **(A)** Outline of biochemical strategy to isolate ribosome protected fragments (RPFs), corresponding to ribosome associated mRNAs. **(B)** Normalized RPF read coverage is shown for ZFP36 mRNA in WT and *Zfp36* KO cells. RPF coverage is lost in *Zfp36* KO cells downstream of the site of gene disruption (Taylor et al., 1996), confirming our biochemical strategy faithfully identifies translating mRNAs. **(C)** Analysis of shifts in ribosome association are shown as in Figure 3B, further stratified by the magnitude of ZFP36 binding. The top 20% of sites (red) show similar shifts to sites overall (violet) for both 3’UTR (top panel) and CDS (bottom panel) in ribosome profiling experiments.

Levels of ribosome-associated mRNA are related to total abundance. To evaluate changes in translational efficiency (ΔTE) in *Zfp36* KO versus WT T-cells, ribosome profiling fold-changes were normalized to those from RNAseq. We then used Gene Set Enrichment Analysis (GSEA) to examine the distribution of ZFP36 targets among mRNAs ranked by ΔTE (Subramanian et al. 2005). ZFP36 3’UTR and, more significantly, CDS binding targets were strongly enriched for increased TE in *Zfp36* KO cells (Figure 3C). In addition, ZFP36 targets with highly robust 3’UTR binding showed more significant effects on TE than ZFP36 3’UTR targets overall. As a striking confirmation of these results, normalized ribosome coverage on robust 3’UTR targets *Tnf* and *Ifng* was significantly higher in *Zfp36* KO cells than WT (Figure 3D), despite no detectable difference in overall mRNA abundance (Figure 2A). Crucially, ribosome coverage averaged across all mRNAs was not appreciably different between KO and WT cells, indicating specific effects on ZFP36 targets (Figure 3E). Notably, the pattern of ribosome association along these and other transcripts is remarkably consistent between WT and *Zfp36 KO* cells, but with altered magnitude. Mechanistically, this observation indicates that ZFP36 prevents association of mRNAs with ribosomes, but does not impact elongation. These results indicate repression of mRNA target translation by ZFP36 during T-cell activation, likely at the level of initiation.

### ZFP36 negatively regulates T-cell activation kinetics

ZFP36 target mRNAs pointed to multilayered control of T-cell function, including its reported regulation of effector cytokines (e.g. *Il2, Ifng*, *Tnf*, *Il4, Il10*; Figure S4A). Novel targets spanned direct components of the TCR complex (e.g. *CD3d*, *CD3e*), co-stimulatory and co-inhibitory molecules (e.g. *Cd28*, *Icos, Ctla4*), signaling (*Fyn*, *Sos1*, *Akt1*), and transcriptional response (e.g. *Fos*, *Nfatc1*, *Nfkb1*). As an unbiased assessment, we examined the distribution of ZFP36 targets in high-resolution gene expression time courses of CD4+ T-cell activation (Yosef et al. 2013). ZFP36 targets were enriched for mRNAs, like its own, that were rapidly induced after T-cell activation, and targets were depleted among mRNAs with stable expression or delayed induction (Figure 4A). Gene Ontology (GO) enrichments spanned many basic metabolic and gene regulatory functions, in addition to signal transduction, cellular proliferation, and apoptosis (Figure 4B; Table S2).

**Figure 4.**
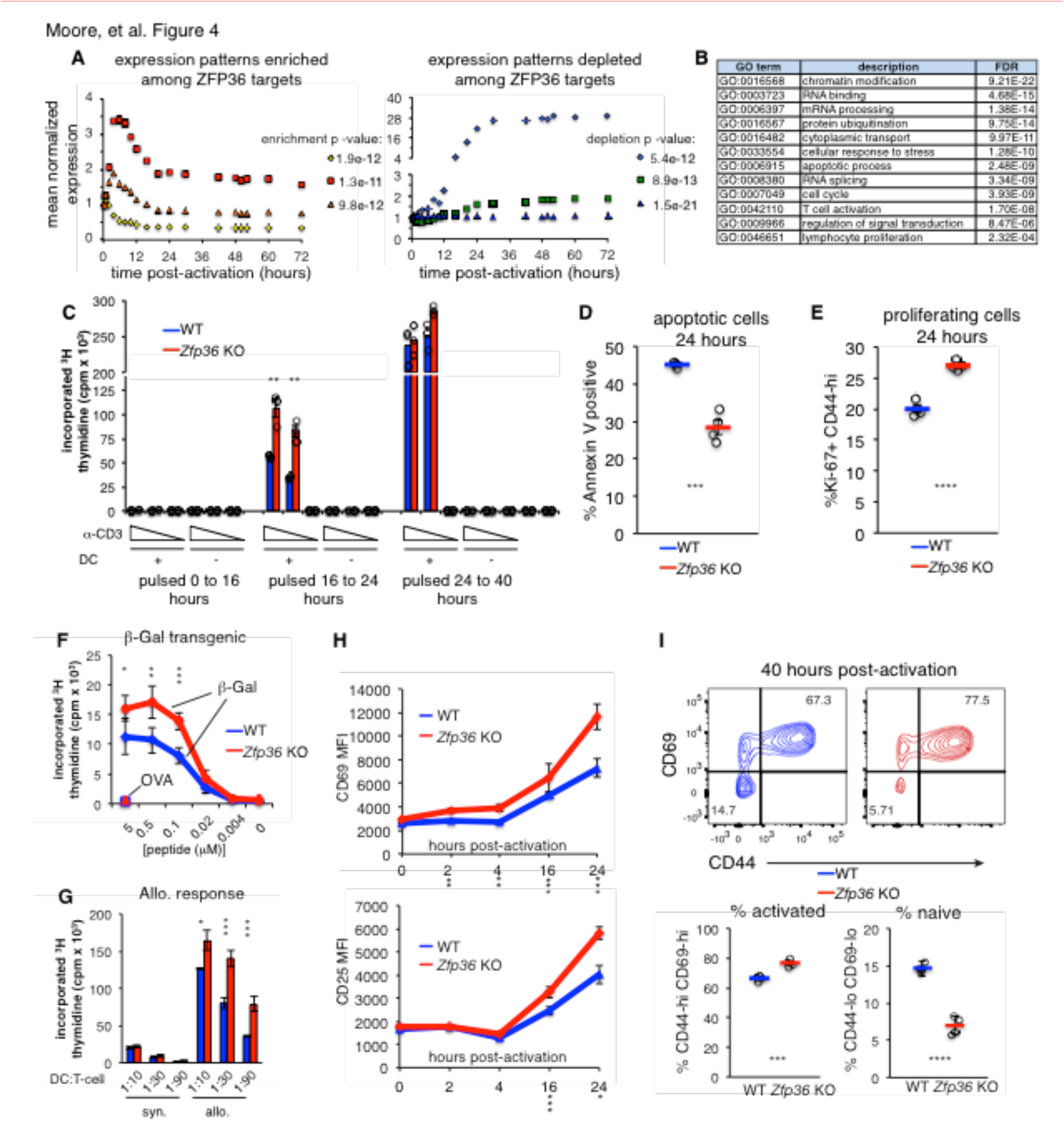
ZFP36 regulates T-cell activation kinetics. **(A)** Gene expression patterns from a T-cell activation time course (Yosef et al. 2013) were partitioned by k-means, and enrichment of ZFP36 3’UTR and CDS targets was determined across clusters (Fisher’s Exact Test). Mean expression of genes in the 3 clusters most enriched (left) or depleted (right) for ZFP36 targets is plotted. **(B)** Enriched GO terms among ZFP36 HITS-CLIP targets (full results in Table S2). **(C)** Proliferation of naïve CD4+ *Zfp36* KO and WT T-cells in the indicated time windows after activation, measured by ^3^H-thymidine incorporation **(D)** Fractions of annexin-V+ and **(E)** Ki67+ CD4+ T-cells 24 hours post-activation. Mean +/- S.E.M. is shown; circles are individual mice (n=3-4 per genotype). **(F)** Proliferation of BG2 TCR-transgenic CD4+ T-cells cultured with DCs pulsed with cognate (β-gal) or irrelevant (OVA) peptide. Mean +/-S.E.M. is shown (n=5 mice per genotype). **(G)** Proliferation of CD4+ T-cells co-cultured with syngeneic (C57BL6/J) or allogeneic (Balb-c) DCs. Mean +/- S.E.M. of three replicate cultures is shown. **(H)** Levels of CD69 and CD25 after activation of *Zfp36* KO and WT naïve CD4+ T-cells. Mean +/- S.E.M. is shown (n=3-4 mice per genotype). **(I)** Naïve and effector subsets 40 hours post-activation in *Zfp36* KO and WT CD4+ T-cells. Representative plots are shown (top), along with mean +/- S.E.M and circles for individual mice (n=4 per genotype). For **(C-I)**, results of two-tailed t-tests: * = p<0.05; ** =p<0.01; *** =p<0.001. See also Figure S4.

**Supplementary Figure S4.**
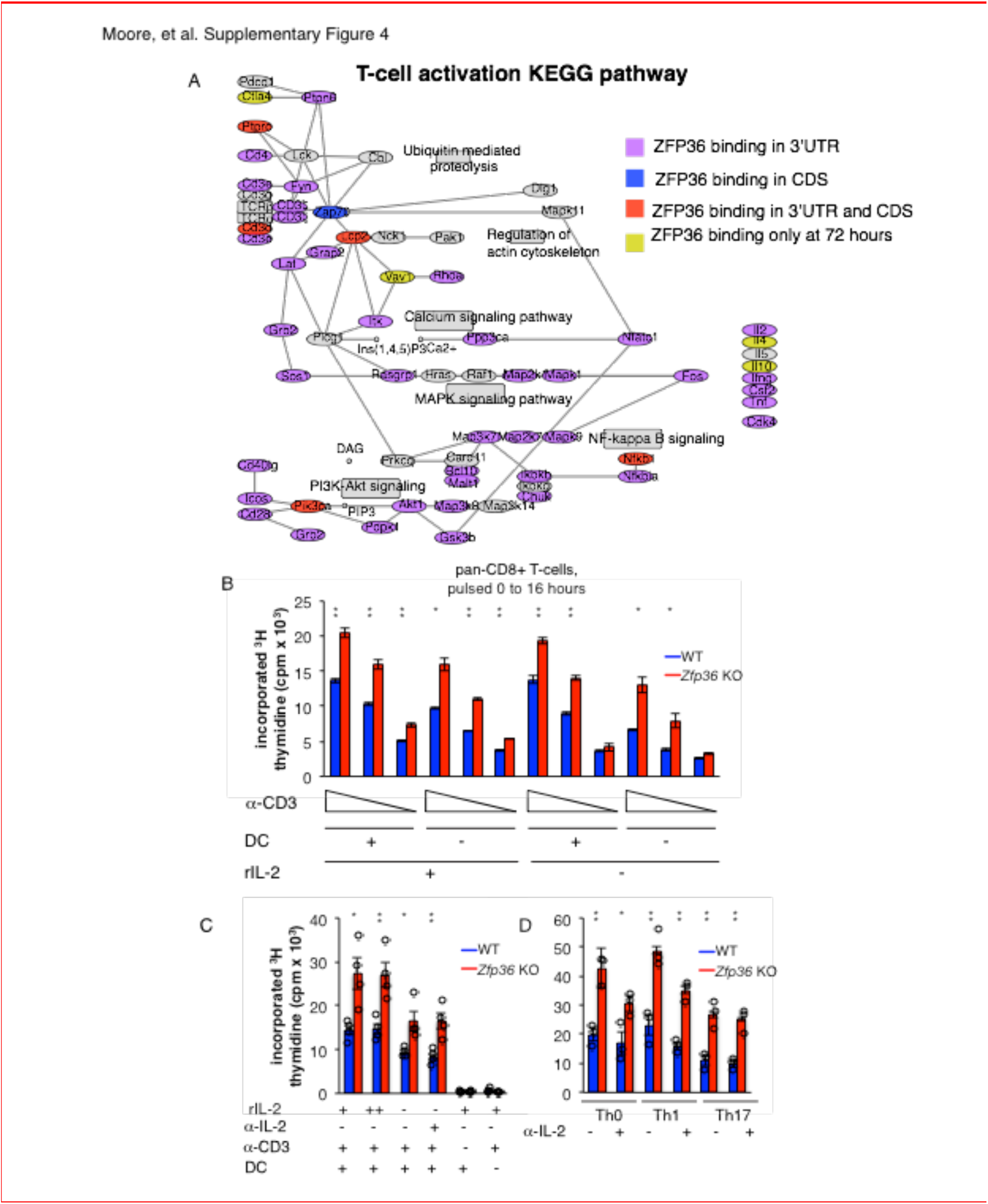
ZFP36 regulates early activation across T-cell lineages. **(A)** The T-cellactivation KEGG pathway is shown with robust ZFP36 CLIP targets shaded based on the location and timing of ZFP36 binding. **(B)** Measurement of proliferation after activation of CD8+ T-cells, varying TCR signal strengths (anti-CD3); co-stimulation (DCs); and the presence of recombinant IL-2. The mean c.p.m. +/ - S.E.M. of three replicate cultures per condition is shown for a representative experiment. **(C)** Measurement of proliferation by thymidine incorporation after activation of naïve CD4+ T-cells is shown in the presence or absence of excess recombinant IL-2; the presence of neutralizing IL-2 antibodies; and **(D)** under various Thskewing conditions. Mean +/- S.E.M. is shown, with circles indicating individual mice (n=3-4 per genotype). For **(B-D)**, results of two-tailed t-tests are shown above relevant comparisons: * = p<0.05; ** =p<0.01; ***=p<0.001.

Functional clustering of ZFP36 targets in proliferation and apoptosis prompted us to investigate potential regulation of T-cell proliferation. In thymidine incorporation assays, naïve *Zfp36* KO CD4+ T-cells showed greater proliferation than WT from 16 to 24 hours post-activation (Figure 4C). Similar results were obtained with CD8+ T-cells (Figure S4B). This increase reflected decreased apoptosis (Figure 4D) and increased numbers of proliferating cells (Figure 4E) in KO versus WT cultures. We examined whether an action on IL-2 might account for enhanced proliferation, as increased IL-2 production in *Zfp36* KO T-cells has been reported (Ogilvie et al. 2005), and our HITS-CLIP data confirmed direct interaction. *Zfp36* KO T-cells proliferated more than WT both in the presence of excess recombinant IL-2 or neutralizing anti-IL-2 antibody, as well as in different Th polarizing conditions, indicating the effect is not solely IL-2-dependent (Figure S4C–D). In summary, T-cells from *Zfp36* KO mice show enhanced proliferation shortly after activation under all conditions examined.

Anti-CD3 is not a physiologic stimulation, so we next examined proliferative responses to MHC-peptide-mediated TCR binding. First, we bred WT and *Zfp36* KO mice with a transgenic, class-II restricted TCR specific for a β-galactosidase-derived antigen (BG2). BG2 transgenic *Zfp36* KO cells showed greater proliferation than WT across a broad titration of cognate peptide, but not irrelevant peptide (Figure 4F). Second, *Zfp36* KO T-cells also showed greater proliferation than WT in response to allogeneic DCs (Figure 4G). Therefore, *Zfp36* KO cells show an exaggerated proliferative response upon MHC-peptide stimulation over a range of signal strengths.

Analysis of canonical T-cell activation markers revealed enhanced induction of CD69 and CD25 in *Zfp36* KO versus WT cells over the first 24 hours post-activation (Figure 4H). At 40 hours post-activation, a greater proportion of *Zfp36* KO versus WT had transitioned from a naïve to effector surface phenotype (Figure 4I). Notably, thymidine incorporation data showed enhanced proliferation of *Zfp36* KO cells early after activation, but similar rates in *Zfp36* KO and WT cells after 24 hours (Figure 2C). Collectively, these results show accelerated activation kinetics in the absence of ZFP36.

### ZFP36 regulation of T-cell activation is cell-intrinsic

The accelerated activation of *Zfp36* KO T-cells could in principle reflect the activity of other cell subsets or inflammatory signals in *Zfp36* KO mice. To test for a T-cell-intrinsic function, we generated mixed bone marrow (BM) chimeras, allowing isolation of WT and KO T-cells that develop in the same in vivo milieu (Figure 5A). Naïve *Zfp36* KO T-cells sorted from chimeras showed greater proliferation than WT 24 hours post-activation, indicating cell-intrinsic effects (Figure 5B). To assess the potential impact of secreted factors, chimera-derived WT and *Zfp36* KO cells were re-mixed 1:1 ex vivo. Here, differences between *Zfp36* KO and WT cells were still significant, but blunted compared to separate cultures. Thus, secreted factors and autocrine signals may contribute to more rapid activation of *Zfp36* KO cells, but do not fully account for it. Interestingly, the reduced proliferation of *Zfp36* KO cells in mixed (Figure 5B, right panel) versus separate (left panel) cultures indicate that WT cells can exert a suppressive effect on KO cells. Thus, accelerated activation in *Zfp36* KO cells may in part reflect compromised autoregulatory and/or suppressive functions. Three days after activation, mixed cultures remained skewed in favor of *Zfp36* KO cells, again confirming accelerated expansion (Figure 5C). These results show that ZFP36 regulation of T-cell activation is cell-intrinsic, and that ZFP36 normally functions to restrain T-cell activation.

**Figure 5.**
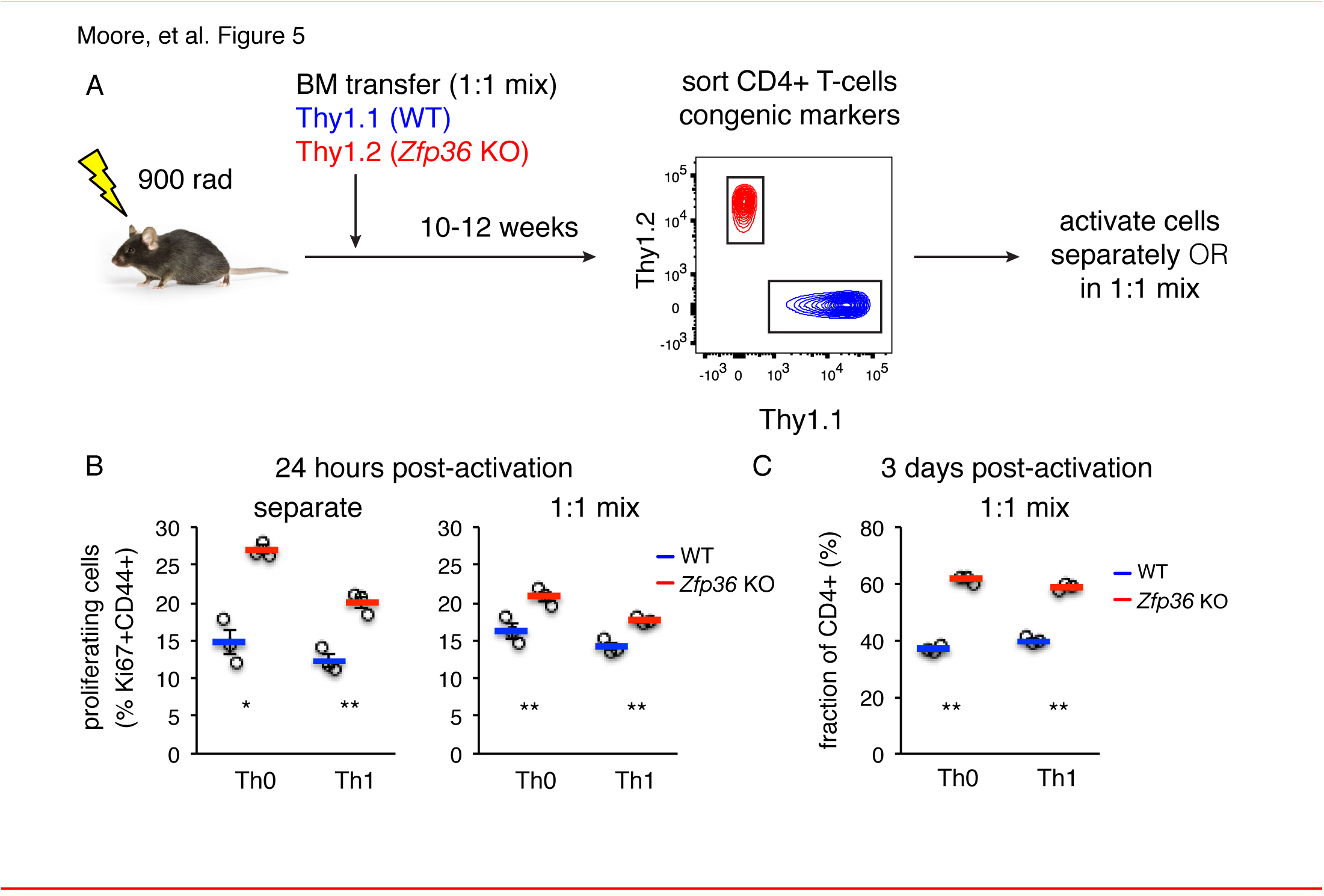
ZFP36 regulation of T-cell activation kinetics cell-intrinsic. **(A)** Lethally irradiated mice were reconstituted with congenically marked WT and *Zfp36* KO BM to generate mixed chimeras. 10-12 weeks after reconstitution, naïve CD4+ WT and *Zfp36* KO T-cells were sorted, then activated ex vivo separately or mixed 1:1. **(B)** Proliferating Ki67+ cells were measured 24 hours after activating naïve CD4+ T-cells under Th0 or Th1 conditions. **(C)** Cultures with a 1:1 starting ratio of naïve WT and *Zfp36* KO CD4+ T-cells were examined 3 days post-activation.

### Downstream effects of ZFP36 regulation

The efficient in vitro responses of *Zfp36* KO T-cells suggest they are functional, but respond with altered kinetics. To examine the downstream consequences of this differential regulation, HITS-CLIP and RNAseq analyses were done in Th1-skewed CD4+ T-cells 3 days after activation. ZFP36 binding site features in cells activated for 3 days were similar to ones identified at 4 hours (Figure S5A–B; Table S3), but results from transcriptome profiling were strikingly different at these two time points (Figure 6A). First, in contrast to subtle effects observed at the 4 hour time point, many transcripts showed highly divergent expression in *Zfp36* KO versus WT T-cells 3 days after activation (Figure 6A). However, these differences were not correlated to ZFP36 HITS-CLIP binding at 72 hours (Figure 6B). Thus, the absence of ZFP36 in the early phases of T-cell activation can lead to significant secondary effects downstream.

**Supplementary Figure S5.**
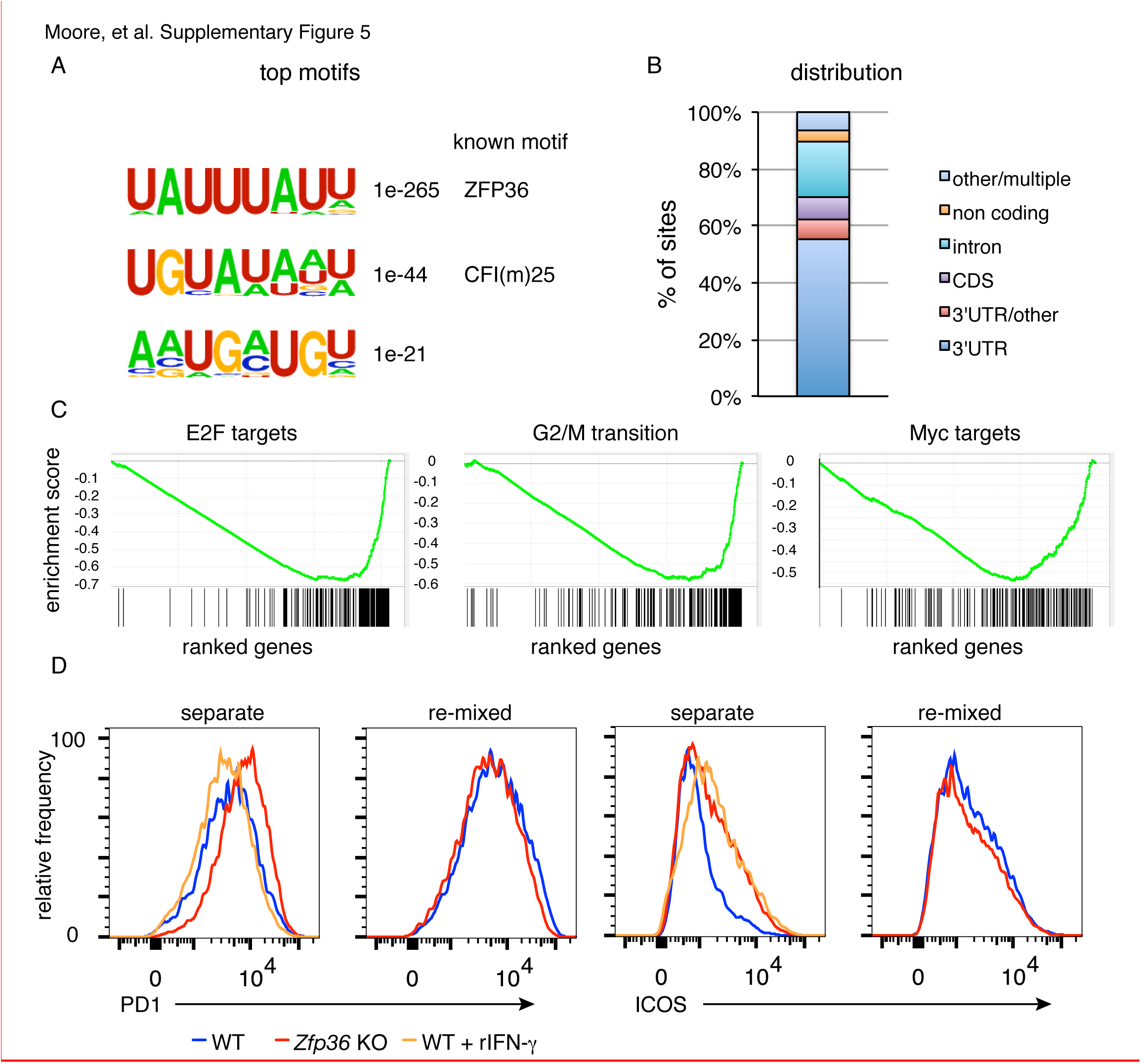
Analysis of ZFP36 function late after T-cell activation. **(A)** Top enriched motifs and **(B)** annotation of ZFP36 HITS-CLIP sites from CD4+ T-cells activated for 3 days under Th1 conditions. Site distribution was similar to the 4 hour time point, except for increased intronic binding, which may reflect increased nuclear permeability in these rapidly cycling cultures. **(C)** GSEA analysis found that genes promoting proliferation and cell division, including transcriptional targets of E2F and Myc, were down-regulated in *Zfp36* KO Th1 cells versus WT 3 days after activation. Enrichment score distributions are plotted, with the distribution of interrogates genes shown below. **(D)** WT and *Zfp36* KO CD4+ T-cells sorted from mixed BM chimeras were analyzed for PD-1 and ICOS expression after long-term (13 days) activation under Th1 conditions. Separated cultures of WT and *Zfp36* KO confirmed differential expression of these receptors, but expression was similar in 1:1 ‘re-mixed’ cultures. Adding recombinant IFN-γ (orange) at 20 ng/ml (similar to levels measured in KO culture supernatants) did not shift PD-1 expression in WT cells toward levels observed in *Zfp36* KO, but did so for ICOS expression.

**Figure 6.**
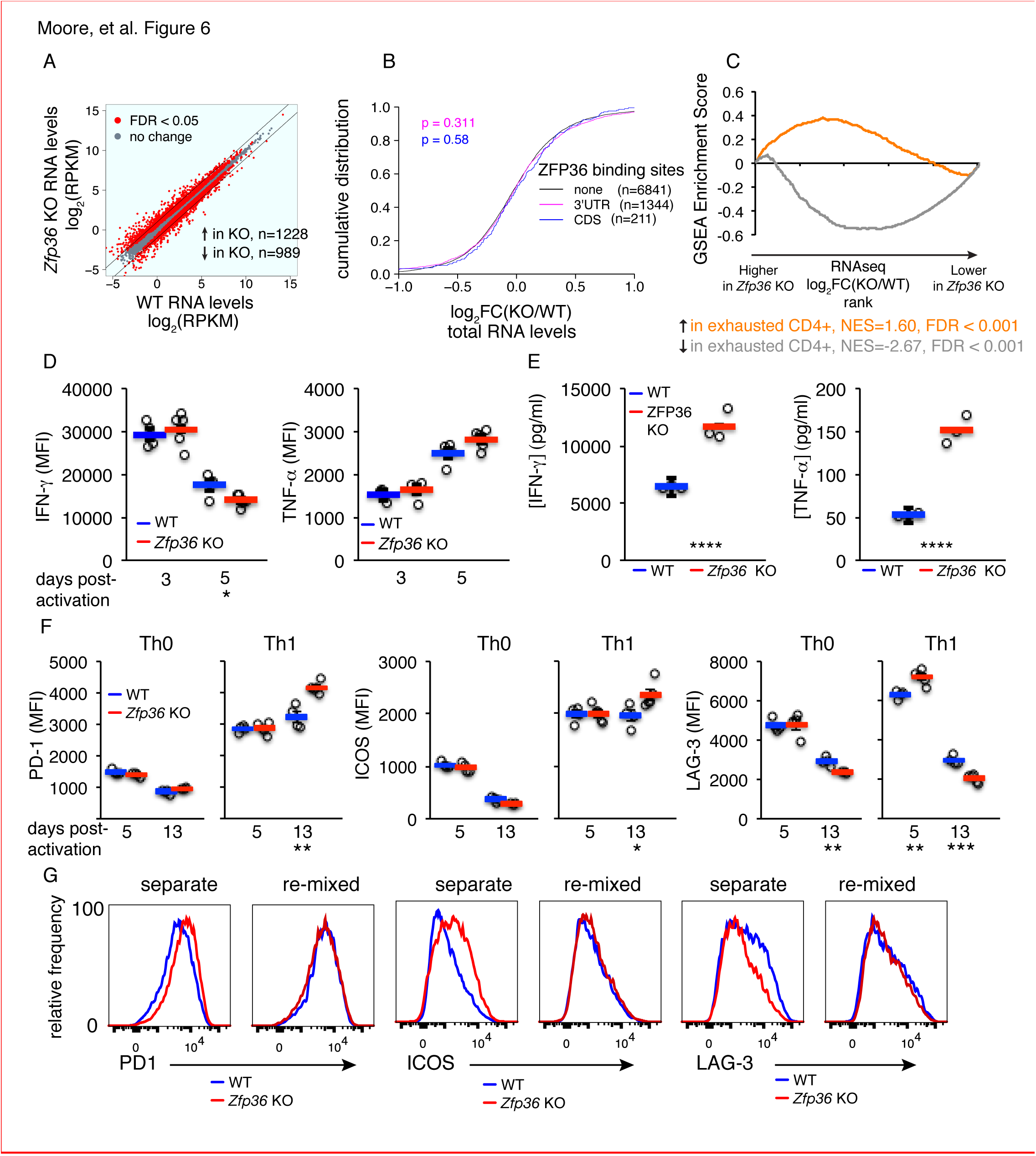
Accelerated signs of in vitro T-cell exhaustion in absence of ZFP36. **(A)** Log2-transformed RPKM values from *Zfp36* KO versus WT CD4+ Th1 cell RNAseq 72 hours post-activation, with red indicating differential expression (FDR < 0.05). Lines mark 2-fold changes. **(B)** Log2-transformed fold-changes (KO/WT) plotted as a CDF, for mRNAs with 3’UTR, CDS, or no significant ZFP36 HITS-CLIP. Numbers of mRNAs in each category (n) and p-values from KS tests are indicated. **(C)** The gene expression profile in *Zfp36* KO CD4+ T-cells 72 hours post-activation was compared to reported profiles of CD4+ T-cell exhaustion using GSEA. Upregulated (orange) and downregulated (gray) gene sets in exhausted T-cells showed strong overlap with corresponding sets from *Zfp36* KO T-cells (FDR<0.001, hypergeometric). (D)IFN-γ and TNF-α measured by ICS 3 and 5 days after activation of naïve CD4+ T-cells. **(E)** IFN-γ and TNF-α in culture supernatants 3 and 5 days after activation of naïve CD4+ T-cells. **(F)** PD-1, ICOS, and LAG-3 expression 5 and 13 days after activation under Th0 or Th1 conditions. **(D-F)** show mean +/- S.E.M.; circles areindividual mice (n=3-5 per genotype). **(G)** Measurements as in **(F)** for *Zfp36* KO and WT CD4+ T-cells derived from mixed BM chimeras. Cells were activated under Th1 conditions for 13 days, either separately or mixed 1:1. Results of two-tailed t-tests: * = p<0.05; ** =p<0.01; *** =p<0.001; **** =p<0.0001. See also Figure S5.

GSEA with RNAseq data from Th1 *Zfp36* KO cells 3 days post-activation showed reduced activity of transcription factors driving proliferation (e.g. Myc and E2F; Figure S5C) and strong overlap with previously described signatures of T-cell exhaustion (Figure 6C; (Crawford et al. 2014)). Consistent with a late activated and exhausted phenotype in *Zfp36* KO cells, the enhanced production of IFN-γ at early time points (Figures 2A and 3D) gave way to comparable production by day 3 and reduced production by day 5 (Figure 6D). Moreover, TNF-α production was not significantly different 3 or 5 days post-activation (Figure 6D), despite large differences early on (Figures 2A and 3D). In summary, relieved translational control drives elevated cytokine production in *Zfp36* KO cells early post-activation, but enhanced production dissipates downstream due to more rapid expansion and exhaustion. Nonetheless, the net result of these shifting dyanmics is higher accumulation of IFN-γ and TNF-α in *Zfp36* KO culture supernatants 72 hours post-activation (Figure 6E).

We examined co-inhibitory and co-stimulatory checkpoint proteins that are linked to T-cell exhaustion, and found elevated expression of PD-1 and ICOS at late time points in *Zfp36* KO cells, and more rapid peaking of LAG-3 (Figure 6F). Interestingly, these effects were observed in Th1 but not Th0 conditions, suggesting a dependence on Th1 cytokines. To test this dependence, and to examine whether elevated receptor expression was T-cell-intrinsic, we analyzed cells derived from mixed BM chimeras. This analysis confirmed differential, T-cell-intrinsic expression of these receptors (Figure 5G). However, re-mixing WT and KO cells ex vivo neutralized these differences, indicating they are driven by secreted factors. We tested whether recombinant IFN-γ, supplemented at levels measured in KO cultures, could cause elevated receptor expression in WT Th1 cells, and found it promoted ICOS but not PD-1 upregulation (Figure S5D). These results indicate that Th1 cytokines, including but not only IFN-γ, can drive an exhaustion-like phenotype. The absence of ZFP36 promoted this phenotype in vitro, due to more rapid activation and expansion coupled with greater accumulation of Th1 cytokines.

### ZFP36 regulates antiviral immunity

The accelerated response of *Zfp36* KO T-cells, and the potential for accelerated exhaustion, led us to examine the effects of ZFP36 regulation of in vivo. We first determined that naïve *Zfp36* KO mice had normal T-cell levels in peripheral blood and no defects in thymocyte development (Figure S6A–B). Total splenocytes, including T-cells, were slightly reduced in *Zfp36* KO versus WT mice (Figure S6C), but proportions of total CD4+ and CD8+ T-cells were normal (Figure S6D). Proportions of naïve and CD25+ nTeg CD4+ T-cells were also normal in KO mice, and naïve CD8+ T-cells were only very slightly decreased. (Figure S6E–F). Stimulation with phorbol myristate acetate (PMA) and ionomycin showed that more CD4+ and CD8+ T-cells produced IFN-γ in KO versus WT splenocytes, but levels of IL-4 and IL17A production were comparable (Figure S6G). Thus, a larger proportion of T-cells are poised for a Th1, IFN-γ producing phenotype in *Zfp36* KO mice in vivo, consistent with observed dysregulation of Th1 cytokines. However, the ability of sorted, naïve CD4+ T-cells to skew to Th1, Th17, and iTreg subsets ex vivo was similar in WT and *Zfp36* KO cells, indicating that loss of ZFP36 does not fundamentally alter the potential for Th cells to differentiate to diverse functional subsets (Figure S6H–I).

**Supplementary Figure S6.**
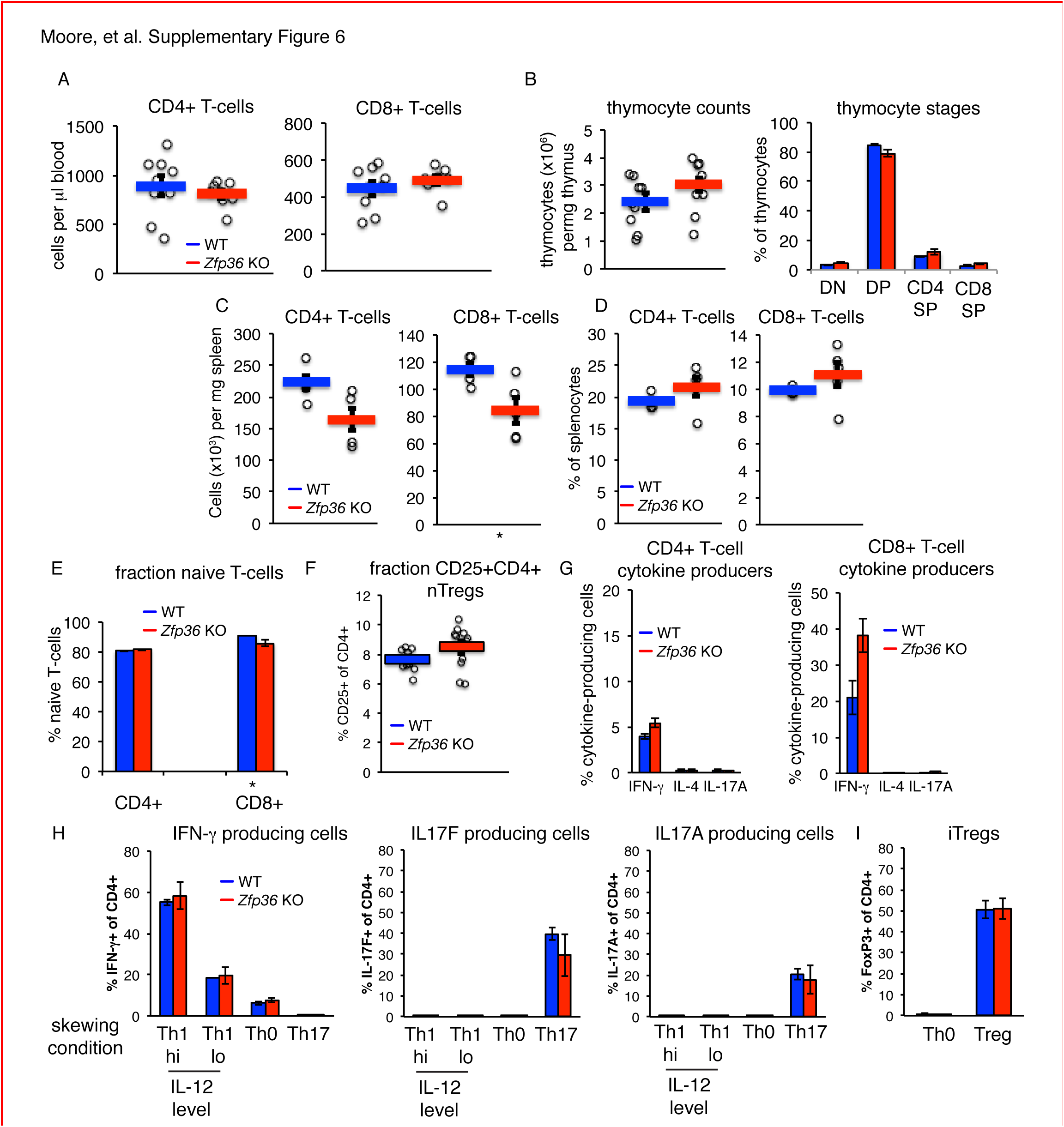
The T-cell compartment in naÏve *Zfp36* KO mice is largely normal. **(A)** Countsof CD4+ and CD8+ T-cells in peripheral blood of WT and *Zfp36* KO mice (n=9-10 per genotype). Mean values +/- S.E.M are shown; circles are individual mice (n=9-10 per genotype) **(B)** Counts of thymocytes and distribution among T-cell development stages in WT and *Zfp36* KO mice (n=11 per genotype; DN=double negative; DP=double positive; CD4-SP=CD4 single positive; CD8-SP=CD8 single positive) **(C)** Counts of CD4+ and CD8+ T-cells in spleen or **(D)** as a proportion of splenocytes in WT and *Zfp36* KO mice (n=5 mice per genotype). **(E)** Percentages of naïve CD4+ and CD8+ T-cells in spleen, defined as CD25-CD62L-hiCD44-lo (n=11 per genotype). For **(A-E),** mean values +/- S.E.M are shown with circles as individual mice. **(F)** Percentages of CD4+CD25+ nTreg cells in spleen (n=11 per genotype). For **(A-F),** mean values +/- S.E.M are shown with circles as individual mice. **(G)** Percentages of CD4+ and CD8+ cells producing the indicated cytokines by intracellular flow cytometry, following 5 hours of PMA/ionomycin stimulation (mean +/- S.E.M. is shown for n=6 per genotype). **(H)** Percentages of CD4+ T-cells producing the indicated lineage-specific effector cytokines is shown under various skewing conditions (mean +/- S.E.M. is shown for n=3 mice per genotype). **(I)** Percentages of FoxP3+ iTreg cells from WT and *Zfp36* KO FoxP3-GFP transgenic mice indicated skewing conditions (mean +/- S.E.M. is shown for n=2 mice per genotype). For **(A-I)** results of two-tailed t-tests are shown beneath relevant panels when significant differences were observed: * = p<0.05. Otherwise, differences were not significant.

The lymphocytic choriomeningitis virus (LCMV) Armstrong strain causes an acute infection leading to massive T-cell expansion and viral clearance in 8-10 days (Dutko and Oldstone 1983). Using MHC-tetramers, we observed accelerated expansion and recession of virus-specific CD4+ (Figure 7A) and CD8+ (Figure 7B) T-cells in *Zfp36* KO versus WT mice in peripheral blood. This result was confirmed in independent experiments focused on early time points post-infection (p.i.), where virus-specific T-cells in *Zfp36* KO mice showed earlier expansion and more rapid upregulation of CD69 (Figure 7C–D). Enumeration of virus-specific T-cells in spleen mirrored dynamics in blood; levels were greater in *Zfp36* KO animals 6 days p.i., but marginally lower by day 10, consistent with more rapid expansion and resolution (Figure 7E–F). Levels of memory T-cells day 40 p.i. were similar in *Zfp36* KO and WT mice. Thus, consistent with data from ex vivo T-cell activation, antigen-specific T-cell response is clinically functional but accelerated in *Zfp36* KO mice during viral infection.

**Figure 7.**
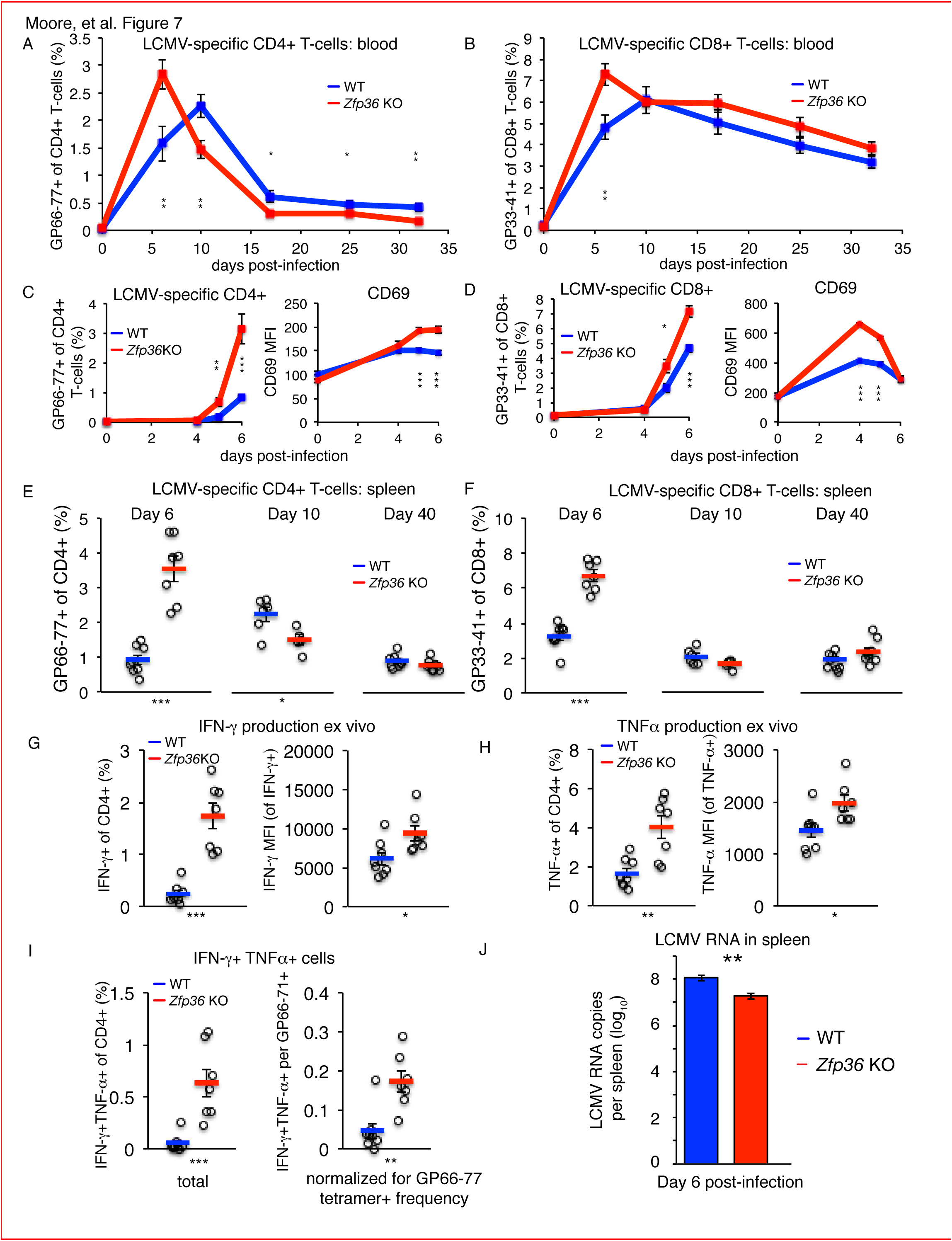
ZFP36 regulates anti-viral immunity. **(A)** Virus-specific CD4+ or **(B)** CD8+ T-cells were tracked in peripheral blood using MHC-tetramers after LCMV Armstrong infection (n=8-9 mice per genotype). **(C)** Virus-specific CD4+ T-cells and CD69 expression on CD4+ T-cells in peripheral blood at early time points post-infection (p.i.) (n=7-8 mice per genotype). **(D)** Virus-specific CD8+ T-cells and CD69 expression on CD8+ T-cells in peripheral blood at early time points p.i. (n=7-8 mice per genotype). **(E)** Virus-specific CD4+ and **(F)** CD8+ T-cells in spleen after LCMV infection (n=5-8 mice per genotype). **(G)** Fraction of CD4+ T-cells producing IFN-γ and TNF-α in splenic CD4+ T-cells 6 days p.i., after ex vivo stimulation with GP66-77 peptide (n=7-8 mice per genotype). **(H)** Levels of IFN-γ and TNF-α (gated on cytokine-producing CD4+ cells) 6 days p.i. after ex vivo stimulation with GP66-77 (n=7-8 mice per genotype). (I)Raw percentage of bifunctional IFN-γTNF-α+ CD4+ cells in spleen 6 days p.i. after ex vivo stimulation with GP66-77 (left), or normalized to percentage of GP66-77 tetramer+ cells (n=7-8 mice per genotype). **(J)** Levels of LCMV genomic RNA in spleen measured by RT-qPCR (n=9-14 per group). For **(A-J)**, mean values +/-S.E.M. are shown, with circles as individual mice. Results of two-tailed t-tests: * = p<0.05; ** =p<0.01; *** =p<0.001; **** =p<0.0001. See also Figures S6 and S7.

Stimulation with LCMV peptides ex vivo revealed higher rates of IFN-γ and TNF-α production in *Zfp36* KO versus WT CD4+ (Figure 7G) and CD8+ T-cells 6 days p.i. (Figure S7A). Numbers of cytokine-producing cells were proportional to LCMV-specific tetramer+ cells (Figure 7E–F). However, levels of IFN-γ and TNF-α protein were significantly greater in CD4+ *Zfp36* KO cytokine-producing cells versus WT (Figures 7H), and TNF-α levels were also higher for CD8+ cells (Figure S7B). In addition, ‘bifunctional’ IFN-γ+TNF-α+ T-cells were more frequent in *Zfp36* KO mice, even when normalized to frequencies of tetramer+ cells (Figures 7I and S7C).

**Supplementary Figure S7.**
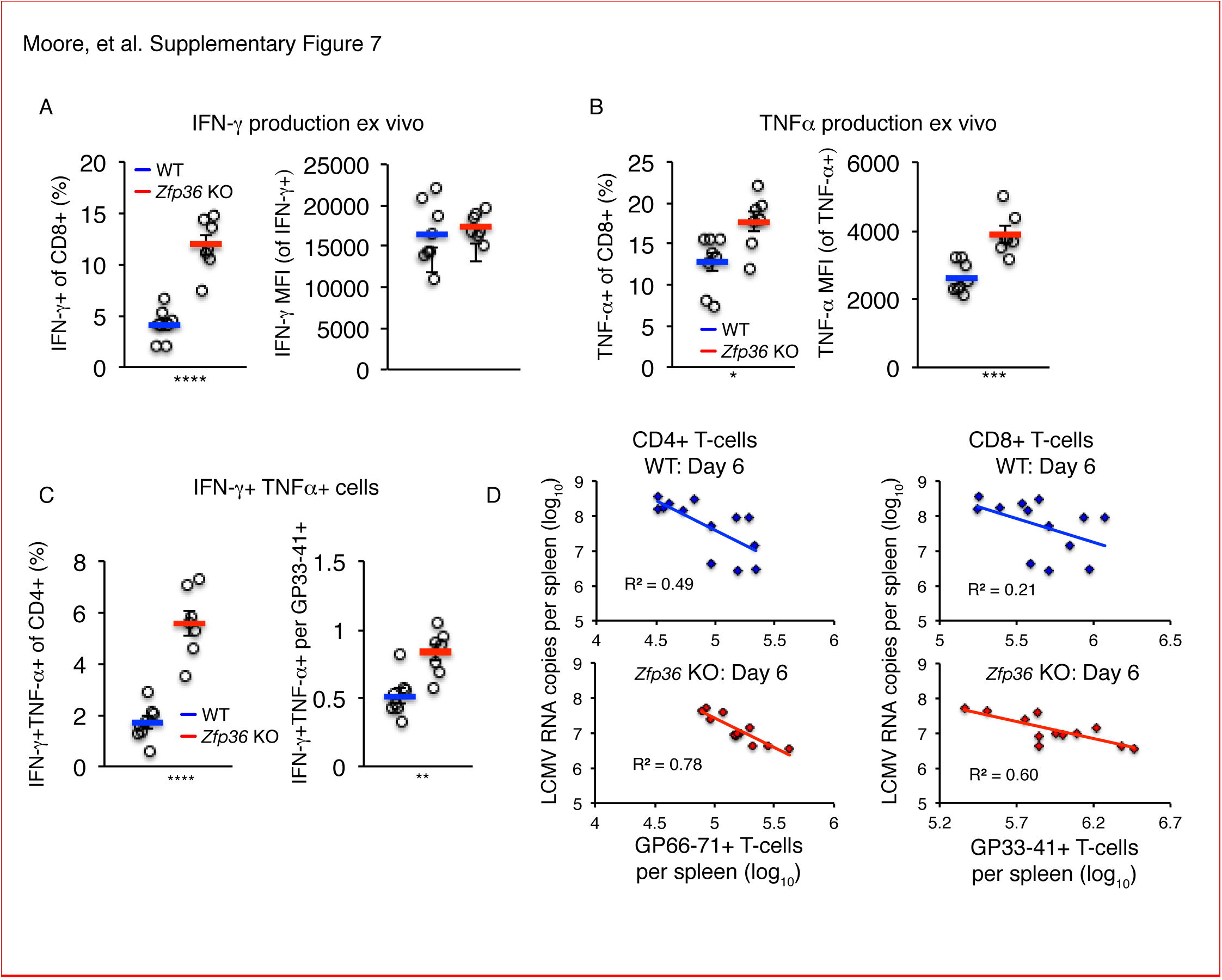
ZFP36 regulates anti-viral immunity. **(A)** The fraction of CD8+ T-cells producingIFN-γ and IFN-γ proteins levels were measured splenic CD8+ T-cells by ICS 6 days post-infection, after ex vivo stimulation with the LCMV antigenic peptide GP33-41 (n=5-7 per genotype). **(B)** The fraction of CD8+ T-cells producing TNF-α and TNF-α proteins levels were measured in splenic CD8+ T-cells by ICS 6 days post-infection, after ex vivo stimulation with the LCMV antigenic peptide GP33-41 (n=5-7 per genotype). **(C)** The raw percentage of bifunctional IFN-γTNF-α+ CD8+ cells in spleen 6 days post-infection after ex vivo stimulation with GP33-41 (left), or normalized to percentage of GP33-41 tetramer+ cells (right) (n=5-7 per genotype). **(D)** Plots depicting the relationship between LCMV load and levels of tetramer+ virus-specific T-cells in spleens of WT and *Zfp36* KO animals, 6 days post-infection. R-squared and p-values are shown for linear regression analysis. For **(A-D),** mean values +/- S.E.M are shown with circles as individual mice. Results of two-tailed t-tests are shown beneath relevant panels when significant differences were observed: * = p<0.05; ** =p<0.01; *** =p<0.001, **** =p<0.0001.

Strikingly, LCMV genomic RNA in spleen was ~10-fold lower day 6 p.i. in *Zfp36* KO versus WT animals, consistent with more rapid clearance of LCMV infection. Viral load correlated inversely with levels of tetramer+ CD4+ and CD8+ T-cells in both *Zfp36* KO and WT mice, consistent with the established role of T-cell response in LCMV clearance (Figure S7D). Collectively, these data demonstrate a remarkable enhancement of anti-viral immunity in the setting of reduced ZFP36 family activity, marked by an accelerated T-cell response and enhanced production of effector cytokines.

## Discussion

Immune response requires rapid, adaptable gene regulation—features uniquely suited to post-transcriptional control. Our studies illuminate a role for ZFP36 RNA binding proteins in controlling the pace of T-cell response, a crucial dimension of adaptive immunity, and tie this to the effectiveness of anti-viral responses in vivo.

Definitive determination of ZFP36 targets in T cells by HITS-CLIP, coupled with transcriptome and ribosome profiling studies, revealed that ZFP36 attenuates T-cell activation by suppressing the abundance and translation of its mRNA targets. The correlation of ZFP36 binding with reduced mRNA abundance is consistent with reports that ZFP36 can destabilize target mRNAs by recruiting degradation factors (Fabian et al. 2013; Lykke-Andersen and Wagner 2005). However, our ability to stratify the relative magnitude of ZFP36 binding using CLIP resolved a more complex trend, with highly robust 3’UTR binding sites (top 20%) showing no detectable correlation with RNA abundance (Figure S2G). This non-uniform trend was observed for 3’UTR but not CDS targets, and affected mRNA abundance but not ribosome association. Importantly, effects on protein levels in the absence of changes in mRNA abundance were confirmed independently for robust 3’UTR binding targets *Tnf*, *Ifng*, and *Cd69*. It is possible that different degrees of ZFP36 association in vivo elicit distinct functional outcomes, through differential RNP localization or downstream effector recruitment. Notably, ZPF36 CLIP showed a broad MW range of ZFP36-RNA complexes with distinct biochemical properties including stability to heat, detergent, and high salt. While the current studies did not uncover distinct mRNA targets across this range, more detailed biochemical studies will be necessary to clarify potentially distinct ZFP36 complexes in vivo, and their potentially distinct roles in different cell types of the immune system.

We further present evidence that ZFP36 suppresses translation of its target mRNAs in T-cells. Endogenous targets and exogenous reporters showed greater ZFP36-dependent suppression of protein versus RNA levels (Figure 2), and ribosome profiling in primary T-cells confirmed direct effects on translation (Tao and Gao 2015; Qi et al. 2012). The strongest effects were linked to a novel class of AREs in coding sequence, uncovered with ZFP36 binding maps. CDS sites correlated with repressed RNA abundance and translation, but a greater level of repression was evident in ribosome association (Figure 3B–D). Intriguingly, some ZFP36 CLIP reads in CDS sites spanned exon-intron boundaries, indicating these associations can form prior to pre-mRNA splicing in the nucleus (Figure S2E). The identification of AREs in the CDS and the possibility of resulting translation control pre-programmed in the nucleus point to novel, unexplored regulatory strategies. Notably, these results differ significantly from iCLIP analyses in macrophages using exogenous GFP-tagged ZFP36, where only 3’-UTR sites correlated with target repression, and only for a ZFP36 construct with mutated MK2 phosphorylation sites (Tiedje et al. 2016). In those studies, the WT ZFP36 construct showed negligible repressive effects, contrasting with our data in 293 and T-cells, and data from other contexts (Tao and Gao 2015; Ogilvie et al. 2009). However, iCLIP data for the transduced WT ZFP36 showed low 3’UTR binding (23%), high intergenic binding (38%), and a preference for GU-rich motifs, diverging sharply from our analysis of endogenous ZFP36 and prior in vivo and in vitro characterizations (Brewer et al. 2004; Worthington et al. 2002). These differences may reflect distinct ZFP36 phosphorylation, and hence regulatory outcomes, or as yet undefined variables related to the different cellular context. A direct comparison is further confounded by the use of exogenous, transduced constructs in macrophage experiments, in contrast to our analysis of endogenous proteins in T cells. Importantly, the methods and reagents reported here allow for systematic analysis of endogenous ZFP36 in future global and cell-type-specific investigations addressing these issues.

The similarity of RNA-binding maps covering both ZFP36 and ZFP36L1 (WT cells) or ZFP36L1 alone (*Zfp36* KO cells) supports redundancy of ZFP36 paralogs, a likely source of robustness in immune regulation. *Zfp36* KO cells are thus likely a partial loss-of-function due to robust ZFP36L1 expression, a notion consistent with the relatively subtle regulatory effects on RNA abundance and translation. Phenotypically, loss of *Zfp36* led to accelerated activation of mature T-cells, but not uncontrolled proliferation or impaired development, which may again may reflect a partial loss of pan ZFP36 activity. Indeed, a prior study reported no effects when *Zfp36l1* was deleted in T-cells, but drastic dysregulation of thymocyte proliferation upon loss of both *Zfp36l1and Zfp36l2* (Hodson et al. 2010). These studies indicate total paralog dosage is critical, but also suggest the importance of the specific balance of ZFP36 paralogs in a defined context. Improved cell profiling methods, such as cell-type-specific tagging of RBPs, may illuminate these complexities in future studies.

The mRNA targets defined by ZFP36 HITS-CLIP span from surface molecules engaged in the earliest steps in T-cell activation to downstream signaling and transcriptional effectors. Targets were strongly enriched for regulation of proliferation and apoptosis, extending prior reports that the ZPF36 family regulates proliferation in early T- and B-cell development and cancer. In each case, the reported mechanism was distinct, spanning regulation of Notch, G1/S phase transition, and Myc, respectively (Galloway et al. 2016; Hodson et al. 2010; Rounbehler et al. 2012). ZFP36 HITS-CLIP identified all of these target pathways in T-cells, consistent with a central function in controlling cell proliferation. However, the phenotype of *Zfp36* KO T-cells is novel and distinct, leading not to uncontrolled proliferation, but to accelerated effector response and resolution. The global effects of ZFP36 repression on RNA abundance and translation were widespread but subtle, and spanned many layers of T-cell function. Functional validation of novel ZFP36 targets, including T-cell activation marker *Cd69* and apoptosis regulator *Bcl2*, suggest factors that likely contribute to this regulation. In addition, mixing experiments indicate a role for both intracellular and secreted factors, and suggest potentially compromised suppressive functions in T-cells lacking ZFP36 (Figure 5B). Thus, ZFP36 regulation of T-cell activation kinetics involves concerted regulation of many biologically coherent targets directing a multifaceted, finely tuned response.

In vivo studies of acute viral infection showed accelerated expansion and recession of virus-specific CD4+ and CD8+ T-cells in *Zfp36* KO animals. *Zfp36* KO T-cells had higher levels TNF-α and IFN-γ protein expression than WT after peptide stimulation, and more ‘bi-functional’ TNF-α/IFN-γ co-producing cells, thought to be important for anti-viral immunity (Crawford et al. 2014). The accelerated activation kinetics and enhanced cytokine production were highly consistent with ex vivo functional experiments and the ZFP36 target profile.

Strikingly, more rapid T-cell expansion coincided with lower accumulated viral titers (or more rapid clearance) in KO animals, indicating ZFP36 regulates anti-viral immunity. Because these studies utilized global *Zfp36* KO animals, it is important to consider the potential roles of other cell types. Mixed BM chimera experiments demonstrated T-cell-intrinsic ZFP36 regulation in vivo. However, viral infection could lead to accelerated T-cell activation in KO animals though non-cell-autonomous mechanisms—for instance, enhanced priming by APCs or exaggerated innate response. Nonetheless, the accelerated T-cell response in *Zfp36* KO animals is likely to contribute to enhanced LCMV immunity, given the central role of T-cells in this process (Matloubian et al. 1994). Accordingly, we observed a quantitative, negative correlation between viral load and antigen-specific T-cell levels in vivo. Importantly, the enhanced anti-viral response in the absence of ZFP36 in mice is accompanied by spontaneous inflammation and autoimmunity that worsen with age. Our results starkly illustrate the delicate balance of protective immunity against destructive inflammation, and reveal post-transcriptional regulation by RBPs as central to this trade-off.

Starting from transcriptome-wide RNA binding maps in T-cells, we uncovered a crucial function for ZFP36 proteins in regulating adaptive immunity. These data suggest carefully titrated inhibition of ZFP36 might serve as a pharmacologic strategy in contexts where accelerated T-cell response to challenge is desirable. Our in vivo LCMV studies demonstrate acute viral infection as one context, but application to other intracellular pathogens warrants investigation. Moreover, the ability to activate T-cells to target tumor antigens and the clinical utility of checkpoint inhibitors raise the possibility of exploring ZFP36 inhibition to enhance tumor immunity. ZPF36 HITS-CLIP identified many targets central to these strategies, including *Cd274* (PD-L1), *Pdcd1l2* (PD-L2), *Icos*, *Cd27*, *Cd28, Ctla-4*, *Btla,* and *Lag3*, suggesting a means for concerted regulation. The autoimmune phenotype of the *Zfp36* KO mouse highlights an important caveat, common to parallel issues seen with clinical use of checkpoint inhibitors. The tools for cell-type-specific analysis of ZFP36, its targets, and its inhibition now exist to investigate and refine this balance.

## Materials and Methods

### Contact for Reagent and Resource Sharing

Information and requests for reagents can be directed to Robert B. Darnell.

### Data Reporting

All data reported is for independent biological replicates. In most cases, one mouse was one biological replicate. For CLIP studies 2-4 littermate mice of the same sex and genotype were pooled for each biological replicate. When performed, technical replicates deriving from the same biological replicate were averaged. For ex vivo studies, including genomic analyses, a sample size of 3-5 biological replicates was judged sufficient based on a power analysis using values from pilot studies, requiring p < 0.05 with 95% power. To account for greater variability, sample sizes were doubled for in vivo studies. Mouse studies were not blinded.

### Mice and cell maintenance

#### Mice

All mouse experiments were approved by The Rockefeller University Institutional Animal Care and Use Committee regulations. *Zfp36* KO mice were a generous gift from P. Blackshear (NIH; (Taylor et al. 1996)). BG2 mice, a TCR transgenic line specific for the class-II-restricted peptide β-gal p726 (NLSVTLPAASHAIPH) from bacterial β-galactosidase, were a generous gift from Nicolas Restifo (NIH; (Tewalt et al. 2009)). Unless otherwise noted, mice of both sexes were analyzed at age 4-6, and comparisons were between littermates.

#### Cell lines

293 TRex cells were maintained under standard conditions (humidified, 37^o^ C, 5% CO_2_) in DMEM supplemented with 10% fetal bovine serum (FBS) and 0.2 mg/ml gentimicin. J55L cells were maintained under standard conditions in RPMI supplemented with 10% FBS, non-essential amino acids, and 0.2 mg/ml gentimicin (R10 media).

#### T-cell cultures

Purified T-cells were cultured with BMDCs at a 30:1 ratio and 0.2 μg/ml α-CD3 (unless otherwise noted) under standard conditions in Iscove’s Minimum Defined Media (IMDM) supplemented with 10% FBS, non-essential amino acids, and 0.2 mg/ml gentimicin. Cytokine conditions, unless otherwise noted, were: Th0 (10 U/ml hIL-2); Th1 (10 U/ml hIL-2 and 5 ng/ml mIL-12); and Th17 (20 ng/ml mIL-6, 10 ng/ml mIL-23, 1 ng/ml hTGF-β). Media with fresh cytokines were replenished every 2-3 days.

#### Bone Marrow-Derived Dendritic Cells

Bone marrow was flushed from tibia, femur, and pelvic bones with 26.5 gauge needles, and RBCs were lysed. Cells were plated at 10^6^ per ml in (IMDM) supplemented with 10% FBS, 0.2 mg/ml gentamycin, and a 1:20 dilution of supernatant from J558L cells stably expressing recombinant GM-CSF. Cells were refreshed on days 3, 5, and 7 with 0.5 ml additional J55L supernatant. At Day 7, non-adherent cells were collected and CD11c+ cells positively isolated with CD11c microbeads (Miltenyi). CD11c+ cells were replated in IMDM with 1:20 J558L supernatant and 50 ng/ml recombinant mouse TNF-α (R&D Systems). At Day 9, non-adherent BMDCs were collected, washed in PBS to remove cytokines, and cryopreserved in a mix of 90% FBS and 10% DMSO.

## Experimental Method Details

### Pan ZFP36 Antisera Generation

Two pan-ZFP36 antisera (RF2046 and RF2047) were produced in rabbits by Covance against the conserved C-terminal peptide APRRLPIFNRISVSE, and successfully confirmed by ELISA, immunoblotting, and immunoprecipitation (IP). Reactivity to paralogs ZFP36, ZFP36L1, and ZFP36L2 was confirmed immunoblotting (Figure S1A) and IP (not shown).

### Plasmid Construction

ZFP36 paralog expression plasmids were generated by PCR-amplification of coding sequences from mouse spleen cDNA, and cloning into the XhoI/NotI sites of the pOZ-N vector (Nakatani and Ogryzko 2003), which has a N-terminal FLAG-HA tag. To generate reporter plasmids, a mammalian codon-optimized, ‘de-stabilized’ Acgfp1 with a PEST degradation targeting sequence from mouse ornithine decarboxylase was ordered as a gBlock (IDT) and cloned in the pcDNA5/FRT/TO vector (Life Technologies) using HindIII and EcoRV sites. The full length 3’UTR of mouse *Ifng*, or a version with the CLIP-defined ZFP36 binding sites deleted, were ordered as gBlocks and subcloned into the Acgfp1 plasmid using EcoRV and NotI sites.

### 293 Transfection assays

Transfection assays used the 293 TREx line (Life Technologies), which expresses the doxycycline-responsive TetR repressor. Cells were maintained under standard conditions with DMEM supplemented with 10% FBS and gentamycin. 293 TREx cells were transfected with Xtremegene9 reagent, using a 3:1 reagent:plasmid ratio and 250 ng total DNA per well in a 24-well dish. 24-hours post-transfection, media was replaced with media containing 100 ng/ml doxycycline to induce reporter expression. Four hours post-induction, cells were harvested an analyzed for GFP expression by FACS on the Miltenyi MACSQuant. DAPI (20 ng/ml) was added to acquisition buffer for dead-cell exclusion.

### T-cell purification

For purification of pan-CD4+ and CD8+ T-cells, splenocytes were cleared of RBCs by hypotonic lysis, and DC populations were depleted with CD11c microbeads (Miltenyi Biotec). T-cells were then purified with CD4 or CD8 microbeads.

CD4+ naïve T-cells were purified by two strategies. In most experiments, CD4+ cells were pre-enriched from pooled splenocytes and lymph nodes (LNs) by positive selection with CD4 microbeads or depletion with CD19, CD11b, and CD8 microbeads. Naïve CD4+CD25-CD44-loCD62L-hi cells were then FACS-sorted to >99% purity. In ribosome profiling experiments and for Western blot time courses (Figure 1A), naïve CD4+ cells were purified to >95% purity with the CD4+CD62L+ isolation kit (Miltenyi).

### T-cell treatment for RNAseq, HITS-CLIP, and ribosome profiling

For RNAseq, HITS-CLIP and ribosome profiling experiments, selective recovery of T-cell RNA was necessary. Thus, BMDCs were fixed prior to co-culture setup in 1% paraformaldehyde (in 1X PBS) for 5 minutes at room temperature, quenched with a 5-fold volume 0.4M lysine prepared in 1X PBS/5% FBS, and washed extensively in PBS/5% FBS. Pilot experiments confirmed that fixed DCs provide co-stimulatory signals to T-cells and induce ZFP36 expression, though at approximately 2-fold lower efficiency than live DCs (not shown). Thus, a 15:1 T-cell:DC ratio was used with fixed DCs. RT-qPCR measurements confirmed that fixation quenched recovery of DC RNA by ~100-fold (not shown), ensuring selective recovery of T-cell RNA. DC-only RNAseq control samples confirmed that recovered reads from DC RNA were negligible (not shown).

### Immunoblotting

Cell lysates were prepared in lysis buffer [1X PBS/1% Igepal/0.5% sodium deoxycholate (DOC)/0.1% SDS supplemented with cOmplete protease inhibitors and Halt phosphatase inhibitors (Roche)]. Protein concentrations were determined by Bradford assay (Biorad), and 10μg total protein per sample were run on NuPAGE gels (Life Technologies) and transferred to fluorescence-compatible PVDF membrane (Millipore). Membranes were blocked in Odyssey PBS-based buffer (LICOR) for 1 hour to overnight, then primary antibodies were added for 1 hour at room temperature. Antibodies used for Western blotting were: TTP (Sigma T5327, 1:500); ZFP36L1 (CST 2119S, 1:500); FUS (Santa Cruz sc-47711, 1:1000; or Novus NB100-562, 1:10000). After 3 washes in 1X PBS/0.05% Tween-20, membranes were incubated with fluorescent secondary antibodies (LICOR, 1:25,000) for 1 hour at room temperature. Membranes were washed 3 times in 1X PBS/0.05% Tween-20, rinsed in 1X PBS, and visualized on the Odyssey Imaging system (LICOR).

### ZFP36 HITS-CLIP

#### Cell Preparation and UV Cross-linking

FACS-sorted CD4+ naïve T-cells were activated as described above in the presence of formalin-fixed DCs. Cells were harvested at 4 hours, or at 72 hours with 2 hours PMA/ionomycin re-stimulation. Harvested cells were UV-irradiated once at 400 mJ/cm^2^ and once at 200 mJ/cm^2^ in ice cold 1X PBS, pelleted, and snap-frozen until use.

##### Bead preparation

ZFP36 antisera or control sera from pre-immune rabbits was conjugated to Protein A Dynabeads (Life Technologies) in binding buffer (0.1 M Na-Phosphate, pH 8.0), and washed 3 times to remove unbound material. IgG was covalently cross-linked to beads with 25 mM dimethylpidilate (DMP) in 0.2 M triethanolamine (pH 8.2) for 45 minutes at room temperature. Beads were washed twice in 0.2 M ethanolamine pH 8.0, then washed several times in PBS/0.02% Tween-20 containing 5X Denhardt’s buffer. Beads were blocked overnight in the final wash prior to use.

##### Lysis and immunoprecipitation

Cell pellets were re-suspended in 250 μl lysis buffer [1X PBS/1% Igepal/0.5% sodium deoxycholate (DOC)/0.1% SDS supplemented with cOmplete protease inhibitors (Roche), 10 mM ribonucleoside vanadyl complexes (RVC)]. 5 μl RQ1 DNAse (Promega) was added and incubated 5 minutes at 37^o^ C with intermittent shaking. For partial digestion, NaOH was added to lysates to 50 mM and incubated at 37^o^ C with shaking for 10 minutes. Alkali was neutralized by addition of equimolar HCl and HEPES pH 7.3 to 10 mM, and SuperRNAsin (Roche) was added to 0.5 U/μl. For over-digested samples, RVC was omitted from lysate preparation, and RNAse A (1:1000, USB) and RNAse I (1:100, Thermo Fisher) were added and incubated at 37^o^ C for 5 minutes. Lysates were cleared by centrifugation (14,000 rpm for 10 minutes), and rocked with beads for 45 minutes to 1 hour at 4^o^ C. Beads were washed:

-Twice lysis buffer containing 5X Denhardt’s Solution
-Twice high detergent buffer (1X PBS/1% Igepal/1% DOC/ 0.2% SDS).
-Twice high-salt buffer (1X PBS/1% Igepal/0.5% DOC / 0.1% SDS, 1M NaCl [final, including PBS])
-Three times low salt buffer (15 mM Tris pH 7.5, 5mM EDTA)
-Twice PNK wash buffer (50 mM Tris pH 7.5, 10 mM MgCl_2_, 0.5% Igepal)

##### Post-IP on-bead processing

Alkaline phosphatase treatment, polynucleotide kinase (PNK) radiolabeling, addition of pre-adenylated 3’ linker, SDS-PAGE, and nitrocellulose transfer and extraction were performed exactly as described (Moore et al. 2014).

##### RNA footprint cloning

CLIP footprints were reverse-transcribed using the Br-dU incorporation and bead-capture strategy, exactly as described (Weyn-Vanhentenryck et al. 2014). Indexed reverse transcription (RT) primers were used (Supplementary Table 5), allowing multiplexing of 8 samples per Miseq (Illumina) run. cDNA was circularized with CircLigase (Epicentre) as described (Weyn-Vanhentenryck et al. 2014), then amplified with PCR primers with Illumina sequencing adapters. Amplification was tracked with SYBR green (Life Technologies) on the iQ5 Real Time Thermocycler (Biorad), and reactions were stopped once signal reached 500-1000 relative fluorescence units (r.f.u.). Products were purified with Ampure XP beads (Beckman) and quantified with the Quant-IT kit (Life Technologies) and/or Tapestation system (Agilent). Multiplexed samples were run on the Illumina Miseq with 75 base pair single-end reads.

#### RNA-seq

Total RNA was extracted Trizol (Life Technologies) and further purified, with on-column DNAse treatment, using HiPure columns (Roche). 500ng to 1 g total input RNA was rRNA-depleted (Ribo-Zero, Illumina), and unstranded, barcoded libraries were prepared with the TruSeq RNA library kit (Illumina). Libraries were run on the Hiseq 2000, obtaining 50 bp paired-end (PE) reads.

#### Ribosome Profiling

##### Cell Preparation

Naïve CD4+ T-cells were purified with the CD4+CD62L+ isolation kit (Miltenyi) and activated in the presence of formalin-fixed DCs, as described above. After 4 hours, 100 μg/ml cycloheximide was added to cultures and briefly incubated at 37^o^ C. Cells were harvested on ice, washed in 1X PBS containing cycloheximide, pelleted, and snap frozen in liquid nitrogen. Four pairs of biological replicates (WT and *Zfp36* KO) were analyzed.

##### Monosome and Ribosome Protected Fragment (RPF) Preparation

Cells were suspended in 0.75 ml polysome lysis buffer [20 mM HEPES pH 7.3, 150 mM NaCl, 0.5% Igepal, 5 mM MgCl_2_, 0.5 mM DTT, cOmplete protease inhibitors, 100 μg/ml cycloheximide, SuperRNAsin (1:1000)]. Lysates were cleared by spinning for 10 minutes at 2,000 rpm to pellet nuclei, followed by 10 minutes at 14,000 rpm for debris. For digestion of polysomes to monosomes, 1 mM CaCl_2_ and 1500 U micrococcal nuclease (MNase, Thermo Fisher) were added to clear lysates and incubated for 45 minutes at room temperature with rocking. Digests were stopped with addition of 5 mM EGTA, then loaded over 10-50% w/w sucrose gradients and spun at 35,000 rpm for 3 hours. Sixteen fractions were collected using the ISCO Density Gradient Fractionation System, tracking monosome elution with UV absorbance at 254 nm.

Fractions containing monosomes and residual disomes were pooled and dialyzed against gradient buffer (20 mM HEPES pH 7.3, 150 mM NaCl, 5 mM MgCl_2_) to remove sucrose. Ribosomes were then dissociated by adding 30 mM EDTA and 0.5 M NaCl, and freed ribosome subunits were pelleted by spinning at 75,000 rpm for 2 hours. The supernatant fraction was extracted once with acid phenol and twice with chloroform, then precipitated with standard ethanol precipitation.

##### RPF Cloning

Pelleted RNA was re-suspended in water, and treated with PNK in the absence of ATP to remove 3’ phosphates. RNA was re-precipitated in ethanol, and 3’ linker addition was performed in 20 μl reactions containing 1 μM pre-adenylated L32 linker (Supplementary Table 5), 400 U truncated RNA ligase 2 (NEB #M0351L), and 10% polyethylene glycol (PEG MW8000) for 2 hours at room temperature. Ligation reactions were resolved on 12.5% denaturing TBE-urea PAGE gels, and RNA fragments in the range from ~50-80 nucleotides were eluted from excised polyacrylamide fragments. RPFs were ethanol-precipitated and cloned using the exact Br-dU incorporation and bead-capture method used for HITS-CLIP (see above) and described elsewhere (Weyn-Vanhentenryck et al. 2014).

#### Flow Cytometry

##### Surface and Intracellular Stains

Flow cytometry data was acquired on the BD LSR Fortessa or LSR II systems. Cell sorting was done on the BD FACSAria. For surface stains, cells incubated with antibodies in FACS buffer (1X PBS/1% FBS) for 20 minutes at 4^o^ C, then washed twice in FACS buffer. For live cell analysis, samples were then re-suspended in FACS buffer containing 20 ng/ml DAPI and acquired directly. For fixed samples, cells were washed twice with PBS after surface staining, then incubate with Live/Dead Fixable Aqua (1:1000 in PBS) for 10 minutes at RT. Cells were washed once with FACS buffer, then fixed and permeabilized using the BD Cytofix/CytoPerm kit. For intracellular stains, cells were then incubated with antibodies diluted in 1X Perm/Wash buffer for 30 minutes at RT, washed twice with Perm/Wash buffer, and acquired.

Unless noted otherwise, flow cytometry data was gated on live (DAPI- or Aqua-negative) single cells, with additional marker gates applied as indicated.

##### Intracellular cytokine staining (ICS)

For ICS, cells were stimulated with PMA (20 ng/ml) and ionomycin (1μM) for 5-6 hours in the presence of GolgiStop (BD) prior to harvesting. Cells were processed and analyzed as described above.

Cytokine production in LCMV studies was measured in response to class-I H-2^d^-restricted GP33-41 (KAVYNFATM) and class-II I-A^b^-restricted GP66-77 (DIYKGVYQFKSV) LCMV peptides. 2 x 10^6^ splenocytes were incubated in 1 ml R10 media (RPMI with 10% FBS, 50 μM β−ME, non-essential amino acids, and gentamycin) with GolgiStop and 0.2 μg/ml class-I peptides or 2μg/ml class-II peptides. After 5-6 hours, cells were harvested and processed as described above. Gates were established using splenocytes from naïve mice and LCMV-infected mice pulsed with irrelevant peptides (class-I OVA p257 [SIINFEKL]; class-II OVA p323 [ISQAVHAAHAEINEAGR]).

##### MHC Tetramers

LCMV GP33*-*41*-*specific H-2D^b^-restricted MHC-tetramer was purchased as a PE conjugate from MBL. APC-conjugated LCMV GP66-77-specific I-A^b^-restricted MHC-tetramer was a kind gift from the NIH Tetramer Core Facility. For class I tetramer, staining was done as described for surface antibody staining using a 1:800 dilution in FACS buffer. For class II tetramer, staining was done at a 1:300 dilution at RT for 1 hour, then cells were washed once in FACS buffer before staining with surface antibodies as described above. In all experiments, tetramer+ cell gated were established using naïve animals and irrelevant, class-matched tetramers as negative controls.

### Thymidine Incorporation assays

200 μl T-cell-DC co-cultures were set up in 96-well plates, and 1 μCi ^3^H-thymidine was added at indicated time points. Cultures were harvested onto glass filter plates (Perkin-Elmer) and dried thoroughly, before addition of 30 μl scintillation fluid per well and acquisition on the Topcount scintillation counter (Perkin-Elmer).

### Apoptosis assays

24 hours after activation, cultures were harvested on ice and stained with surface markers. Prior to acquisition, buffer was removed from samples and Annexin-V-staining buffer containing Annexin-V-PE (BD Biosciences, 2 μl per samples) and 20 ng/ml DAPI was added.

### Bone Marrow Chimeras

Host CD45.1 mice (Jackson Laboratory 002014) were given a lethal dose of gamma radiation (900 rads), then intravenously injected with a 1:1 mix of Thy1.1 WT and Thy1.2 *Zfp36* KO BM cells (4-5x10^6^ total cells). Chimeras were analyzed 10-12 weeks after re-constitution.

### LCMV studies

Mice were infected with LCMV Armstrong strain by intraperitoneal injection of 1 x 10^5^ plaque forming units. At indicated time points, peripheral blood was collected by retro-orbital bleeding and analyzed for virus-specific T-cells using MHC-tetramers (see above). At 6, 10, and 40 days p.i., spleens were analyzed for LCMV-specific T-cells with MHC-tetramers and analysis of cytokine production in response to LCMV peptide antigens (see above). LCMV genomic RNA was quantified in total spleen RNA by RT-qPCR. To determine copy number, a standard curve was generated using a gBlock comprising the LCMV PCR amplicon. LCMV copy number per spleen was calculated using a linear regression of the standard curve, and scaling by appropriate dilution factors.

## Quantification and Statistical Analysis

Details for statistical analysis appear in figure legends. Comparisons between experimental groups were done with two-sided student’s t-tests, with p<0.05 considered significant. Statistics for bioinformatic analyses are detailed below.

## Bioinformatics

### HITS-CLIP

Processing and alignment of HITS-CLIP read data for Br-dU CLIP was done as described ((Shah et al. 2016; Moore et al. 2014; Weyn-Vanhentenryck et al. 2014)). Peak calling was done with the CLIP Toolkit (CTK) command tag2peak.pl, requiring enrichment over background with FDR < 0.01 (Shah et al. 2016). For background determination, a genic model was used including known mouse transcripts extended 10 kb downstream to include non-annotated 3’UTR variants. For analyses in Figure 1E–G and Supplementary Figure 2C, peaks were defined and analyzed separately for 5 WT (Supplementary Table S1A) and 3 *Zfp36* KO biological replicates (Supplementary Table S1B). In these analysis, bindings sites were defined as peak height [PH]>5, from ≥3 biological replicates, with two different antisera. Subsequently, the 5 WT and 3 KO replicates (8 total) were pooled for peak definition to increase depth and sensitivity (Supplementary Table 1D). For these analysis, peaks had to be supported by two different antisera and >5 biological replicates, ensuring support from both WT and KO datasets. Cross-link-induced truncations (CITS) were also determined as described, pooling all biological replicates from all time points ((Shah et al. 2016); Supplementary Table 1E).

Motif enrichment analysis was done on a 30 nucleotide window surrounding peaks supported in at least 3 biological replicates in the indicated dataset. HOMER (Heinz et al. 2010) was used with commands similar to: findMotifsGenome.pl peak_file.txt mm10 output_folder-rna-len 8 Site annotation was done with custom R scripts using the GenomicRanges and TxDb.Mmusculus.UCSC.mm10.ensGene (Transcript Database for mm10).

T-cell activation time course data was downloaded from GEO, and genes were clustered by expression patterns using k means partitioning in Cluster 3.0 (de Hoon et al. 2004). An optimal k of 20, as previously determined (Yosef et al. 2013), was re-confirmed by visualizing a wide range off k values in Java Treeview (Saldanha 2004). The distribution of ZFP36 3’UTR and CDS CLIP targets among k clustered was evaluated with Fisher’s exact test. Average expression values of genes in the 3 most enriched and depleted clusters were plotted. Gene Ontology enrichments were determined with the TopGO package in R (Alexa and Rahnefuhrer, 2016).

### RNAseq

RNAseq reads were aligned against the mouse reference genome (mm10) with STAR with default settings (Dobin et al. 2013). Feature counts per gene were obtained with HTSeq (Anders et al. 2015). Differential gene expression was analyzed with edgeR using a standard t-test pipeline (Robinson et al. 2010). For CDF analysis (e.g. Figure 1H), log2-fold-change values derived from edgeR were plotted. Analysis included genes with average RPKM > 3 in WT or *Zfp36* KO biological replicates in RNAseq, and with sufficient coverage in ribosome profiling (see below) to permit quantification. Genes were classified based on the presence of robust ZFP36 HITS-CLIP peaks (FDR<0.01, supported by >5 biological replicates) in 3’UTR or CDS. Non-exclusive site annotation was used here, meaning some genes have both 3’UTR and CDS sites. Analysis was repeated with exclusive annotation, with similar results (not shown). The negative control set was defined as genes without significant ZFP36 binding in any transcript region. Differences between gene sets were evaluated with a two-tailed Kolmogorov-Smirnov test.

The gene expression profile observed 72 hours after activation of *Zfp36* KO cells was analyzed for significantly overlapping gene sets using GSEA and the Molecular Signature Database (Subramanian et al. 2005). Overlap with published profiles of CD4+ T-cell exhaustion (Crawford et al. 2014) was determined with a GSEA pre-ranked analysis.

### Ribosome Profiling

Initial filtering and processing of ribosome profiling reads was done as for HITS-CLIP. Reads were aligned against a mouse reference transcriptome (Ensembl) using Bowtie 2, and read counts per transcript were calculated (Langmead and Salzberg 2012). Differential expression analysis was done with edgeR using a paired general linear model (GLM). Only genes with cpm (counts per million read) values > 1 in at least 2 biological replicates of any genotype were included. CDF analyses were done as described for RNAseq. The change in translation efficiency (ΔTE) between *Zfp36* KO and WT cells was calculated for each mRNA as the difference in log2-fold-change values (KO/WT) from ribosome profiling and RNAseq. mRNAs were ranked on the ΔTE metric, and distribution of ZFP36 CLIP targets therein was evaluated with a pre-ranked GSEA analysis.

Ribosome profiling read coverage from pooled biological replicates of each genotype was calculated across individual mRNAs with a sliding 20 nt window, normalizing for dataset read depth. Differences between WT and *Zfp36* KO coverage were evaluated with a binomial test. For the metagene coverage plot, all transcripts with at least 10 reads in the region of interest were included and the proportion of reads which centered at each nucleotide position was calculated. The mean coverage for each position is shown relative to the central coding region with additional normalization included for the number of transcripts represented at each position.

## Data Availability

High throughput sequencing data from this study is available at the NCBI Gene Expression Omnibus (GEO) website under accession GSE96076.

## Author Contributions

M.J.M, N.E.B. and R.B.D. conceived and designed the studies. M.J.M., N.E.B., J.J.F., S.P., and I.Z-.S performed all experiments. B.M. and A.Y.R. contributed to design, execution, and interpretation of LCMV studies. K.S. contributed to design and analysis of ribosome profiling experiments. M.J.M. and C.Y.P. analyzed bioinformatics data. M.J.M and R.B.D. wrote the manuscript, with input from all authors.

## Acknowledgements

We thank Y. Yuan and other Darnell lab members for their support and insightful comments. We are indebted to S. Mazel and the Rockefeller FCRC staff for cell sorting and flow cytometry expertise. M.J.M. was supported by a fellowship from the Jane Coffin Childs Fund for Medical Research. This work was supported by grants to R.B.D. from the National Institutes of Health (NS034389, NS081706, R35NS097404), and the Starr Cancer Consortium. R.B.D. is an Investigator of the Howard Hughes Medical Institute.

